# Whole Genome Sequencing of SARS-CoV-2: Adapting Illumina Protocols for Quick and Accurate Outbreak Investigation During a Pandemic

**DOI:** 10.1101/2020.06.10.144212

**Authors:** Sureshnee Pillay, Jennifer Giandhari, Houriiyah Tegally, Eduan Wilkinson, Benjamin Chimukangara, Richard Lessells, Yunus Moosa, Inbal Gazy, Maryam Fish, Lavanya Singh, Khulekani Sedwell Khanyile, Vagner Fonseca, Marta Giovanetti, Luiz Carols Alcantara, Tulio de Oliveira

## Abstract

The COVID-19 pandemic spread very fast around the world. A few days after the first detected case in South Africa, an infection started a large hospital outbreak in Durban, KwaZulu-Natal. Phylogenetic analysis of SARS-CoV-2 genomes can be used to trace the path of transmission within a hospital. It can also identify the source of the outbreak and provide lessons to improve infection prevention and control strategies. In this manuscript, we outline the obstacles we encountered in order to genotype SARS-CoV-2 in real-time during an urgent outbreak investigation. In this process, we encountered problems with the length of the original genotyping protocol, reagent stockout and sample degradation and storage. However, we managed to set up three different library preparation methods for sequencing in Illumina. We also managed to decrease the hands on library preparation time from twelve to three hours, which allowed us to complete the outbreak investigation in just a few weeks. We also fine-tuned a simple bioinformatics workflow for the assembly of high-quality genomes in real-time. In order to allow other laboratories to learn from our experience, we released all of the library preparation and bioinformatics protocols publicly and distributed them to other laboratories of the South African Network for Genomics Surveillance (SANGS) consortium.

## Introduction

In late December 2019, a mysterious viral pneumonia which had infected a number of people in Wuhan, China, was attributed to a new coronavirus [1]. This virus was labelled severe acute respiratory syndrome coronavirus 2 (SARS-CoV-2) and defined as the causal agent of Coronavirus Disease 2019 (COVID-19) [2]. Despite widespread attempts to contain the virus in China, within a few months the outbreak had reached and affected 215 countries and territories around the world, including all countries in Africa.

Localized outbreaks are common when SARS-CoV-2 is introduced in a new geographic area. These outbreaks require urgent testing and tracing, identification of the causative agent and epidemiological investigations to allow for appropriate infection control and for clusters of patients to be identified [3]. Genomic sequencing can be used to rapidly and accurately identify the transmission routes of the pathogen [4,5]. This can then be used to trace the path of transmission within a population and to possibly identify the probable source, potentially leading to an improved public health response [6–12].

On 5 March 2020, the first COVID-19 case was reported in KwaZulu-Natal, South Africa from a traveller from Italy. On 9 March 2020, another returned traveller from Europe started a large chain of infection in a hospital in Durban, KwaZulu-Natal [13]. On 15 of March 2020, South Africa declared a state of emergency and on 27 March a country wide lockdown was implemented, which included grounding most of the international flights.

In this study, we outline the processes that we went through to set up genomic sequencing in order to investigate the previously mentioned hospital outbreak in South Africa. The process was more complicated than we expected as, in addition to needing to generate data quickly, we also encountered problems with reagent stockout, importing sequencing reagents and sample degradation and storage.

## Results

### Sample characteristics

We received 108 COVID-19 positive samples (sampled from the end of March to the beginning of May 2020). Of these, 77 were from nasopharyngeal and oropharyngeal swabs and 31 were extracted RNA. The average age of the patients was 51 years [ranging from 23-91 years], with a gender distribution of 70.5% females and 29.5% males. Of the 108 individuals, 63 were from the hospital outbreak, 30 were from randomly selected positive individuals sampled in the same city but unrelated to the outbreak at the time of sampling. These last samples were used as control for the outbreak investigation. Furthermore, we also sequenced 13 samples delivered from the hospital outbreak but we could not identify the patient or health care worker from the sample ID or the sample ID has been lost in the transport or in storage before coming to our labs.

### SARS-CoV-2 sequencing with limited reagents and stockout during a pandemic in a country in lockdown

We started sequencing the day after we got involved in the outbreak investigation, on 5 April 2020, so there was no time to prepare. Priority was given to genome sequencing rather than qPCR as there were limited sample volumes and results were required urgently for outbreak investigations. One of the 108 samples failed library preparation; the remaining 107 samples were sequenced and used in the final analysis. It is important to note that these samples did not arrive at the same time. There were 12 shipments and the quality of the samples was not optimal as some were sent to multiple laboratories for diagnostics and were not well stored, i.e. many were stored for days or weeks at room temperature. Furthermore, an undisclosed number of samples were heat inactivated, sometimes more than once.

Our laboratory received training in Oxford Nanopore Technologies SARS-CoV-2 sequencing from colleagues from Brazil and our first two sequencing runs involved 24 samples (n=12 each run) that were sequenced on the 7 April 2020. However, our flow cells were over two years old and had less than 600 active pores each, which produced low quality and lower coverage genome (data not shown). We then moved to use the ARTIC protocol [14] on the Illumina Miseq sequencing platform on 8 April 2020.

We starting using the ARTIC protocol with no modification, which included the TrueSeq library preparation step. However, this was a very lengthy and laborious process, which took close to twelve hours of hands on time to produce the libraries for sequencing. Furthermore, we had reagents for only 24 samples as our order of Illumina TruSeq DNA Library preparation kit (x 96 sample libraries), which was placed in February and was enough to produce 480 genomes, had not arrived due to the restrictions on international flights. In order to complete the outbreak investigation, we needed to improvise and look for other reagents that could replace this library preparation kit and for reagents that were already available in South Africa.

We started by approaching colleagues from six genomics laboratories in South Africa, but unfortunately nobody had a suitable library kit that they could lend us. We also phoned many companies in South Africa to see if they had any local stock. We found two NEB Next Ultra II library kits in stock at a local company, Inqaba Biotech. Inqaba Biotech couriered the kit to us from Johannesburg by car on 10 April. We were happy to have a suitable library kit but quickly realized that one of the components, the NEBNext Multiplex Oligo, which was necessary to run the NEB Next Ultra II library kits, was missing. To overcome this, we changed the protocol to perform adapter ligation using remaining Illumina TrueSeq Single DNA Indexes Set A and Set B. This process also involved adapting PCR conditions and skipping some steps of the original NEBNext Ultra II library protocol. Fortunately, our efforts worked and we had good quality fragment sizes, with expected fragments between 300-600bp in size.

We also approached Illumina directly to chase up our TrueSeq order. However, Illumina informed us that they had recently changed distributors in South Africa and that could not fulfil the previous order. At the time of writing this manuscript, 12 weeks after the order, the TrueSeq reagents have still not arrived. However, via the new distributors in South Africa, Separations Pty, Illumina has provided us with two large complementary library kits, the TruSeq and the Nextera Flex. The Nextera Flex library preparation kit was suggested by their technical team as a potentially better and quicker solution to produce SARS-CoV-2 genomes, which we found to be true.

Normally, one produces a qPCR report before attempting sequencing. However, due to the limited RNA and urgent need to generate data to solve the nosocomial outbreak, we generated a qPCR in parallel to the sequencing process and only after we had generated genomes. In summary, the process in the lab involved three days to complete one round of sequencing, qPCR and analysis of the data (Figure 1). The sequencing process involves RNA extraction, RT-PCR, PCR amplification, library preparation, Illumina sequencing, followed by genome assembly and sequence analysis. We started running 12 samples per round, which was increased to 24 per round during the process. In total, we sequenced 108 COVID-19 positive individuals in less than three weeks.

**Figure 1:**
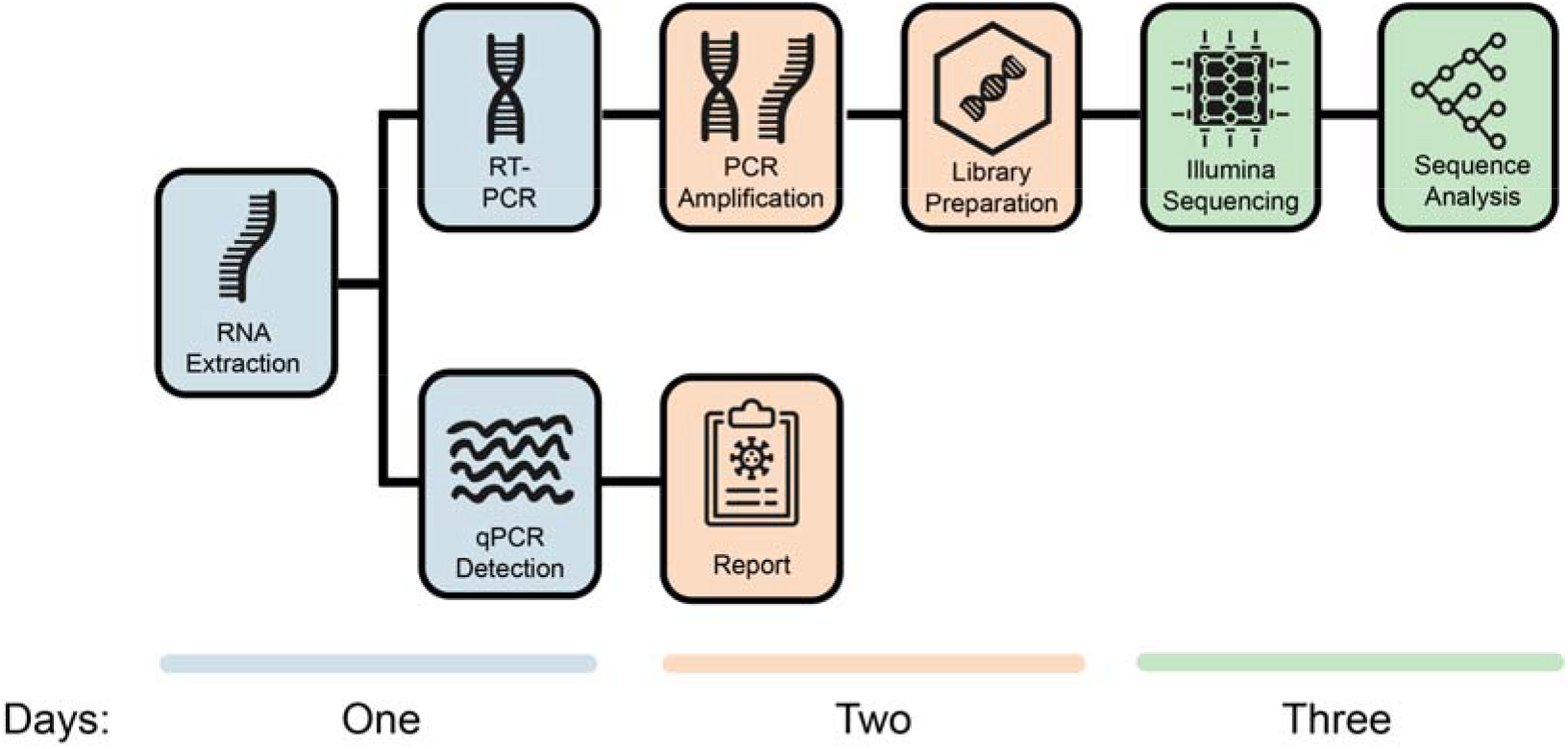
Processes to generate SARS-CoV-2 genomes and qPCR diagnostics in our laboratory. The figure also shows the number of days needed by two senior scientists to generate 24 whole genomes.

In total, 102 of 108 (94.4%) samples had remaining RNA for qPCR. Of the 102 samples, 97 had positive Ct values for all three target genes, and five had at least one undetermined target gene. The median Ct value for the three genes targeted was 24.9 (21.4 – 29.8), with good agreement between the different probes with mean Ct values of 26.513 (22.242 – 31.179), 25.2975 (21.433 – 29.933) and 24.019 (20.853–30.263) for ORF1 ab, S protein and N protein, respectively.

### SARS-CoV-2 Genome Assembly

The generation of high-quality genomes from the sequencing data was generally performed using a 3-step assembly and clean-up workflow (Figure 2). This process involved a last manual step, when whole-genomes are polished and all of the mutations are visually checked from bam alignments for confirmation. We noticed that some of the adaptors were not well filtered in the assembled process and resulted in mis-called mutations. However, this was easy to spot in the bam files and we checked all the mutations visually. On average, our consensus started with 7 (4 - 13) mutations and ended up with 5 (4 - 6). The quality of our consensus sequences clearly increased as indicated by significantly higher (t-test) coverage, concordance, identity and matches after Step 2, and significant increases in matches after Step 3 (Supplementary Figure S1, Table S2). This process produced 107 SARS-CoV-2 genomes, 54 of which were whole genomes (>90% coverage) and 53 partial genomes (mean genome length: 60.5%, IQR 51.5% - 72.4%).

**Figure 2:**
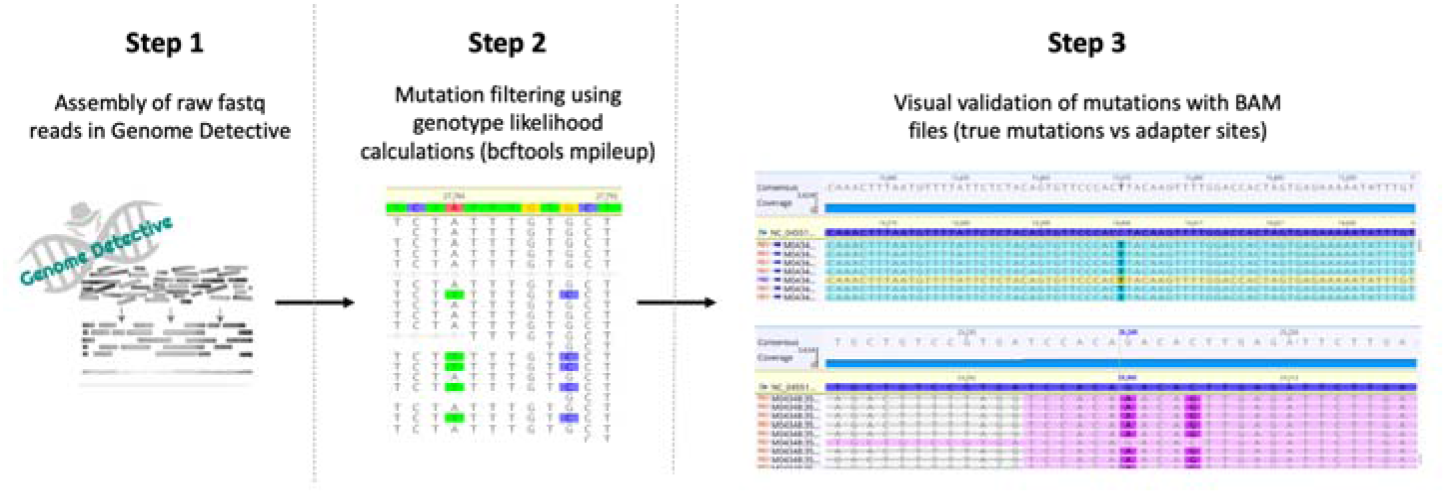
3-step workflow for generation of high-quality genomes. Step 1: Raw reads from Illumina and Nanopore sequencing were assembled using the web-based Genome Detective 1.126 (https://www.genomedetective.com/) platform and its coronavirus typing tool; Step 2: The initial assembly obtained from Genome Detective was polished by aligning mapped reads to the references and filtering out mutations with low genotype likelihoods using bcftools 1.7-2 mpileup method. This calculation determines the probability of a genotype at sites containing reads with various bases (e.g. the probability that position 27784 is A vs T in illustration above); Step 3: All mutations were validated visually with BAM files viewed in Geneious software to ensure that called mutations were true and not part of lingering adapter sites.

### Association between Ct Score and genome length

There was a marked increase in coverage seen in samples with a lower Ct score (Figure 3). There was a clear trade-off between Ct score and genome coverage and we chose three cut-off values to analyze the data, Ct <25, <27 and <30 (Figure 3). We consistently show genomes of significantly higher coverage generated from samples with mean Ct scores below the chosen thresholds. For example, we had Ct score values for 51 of the 54 near fulllength genomes with a mean Ct score of 22.2 (19.7 - 24.2 For this outbreak, we demonstrate that a Ct < 27 show a good trade-off between high-quality near complete genomes Ct score. However, 12 samples with a Ct < 27 did not produce whole genomes (Figure 3). We associated this to sample degradation as many of the samples have been stored at room temperature.

**Figure 3.**
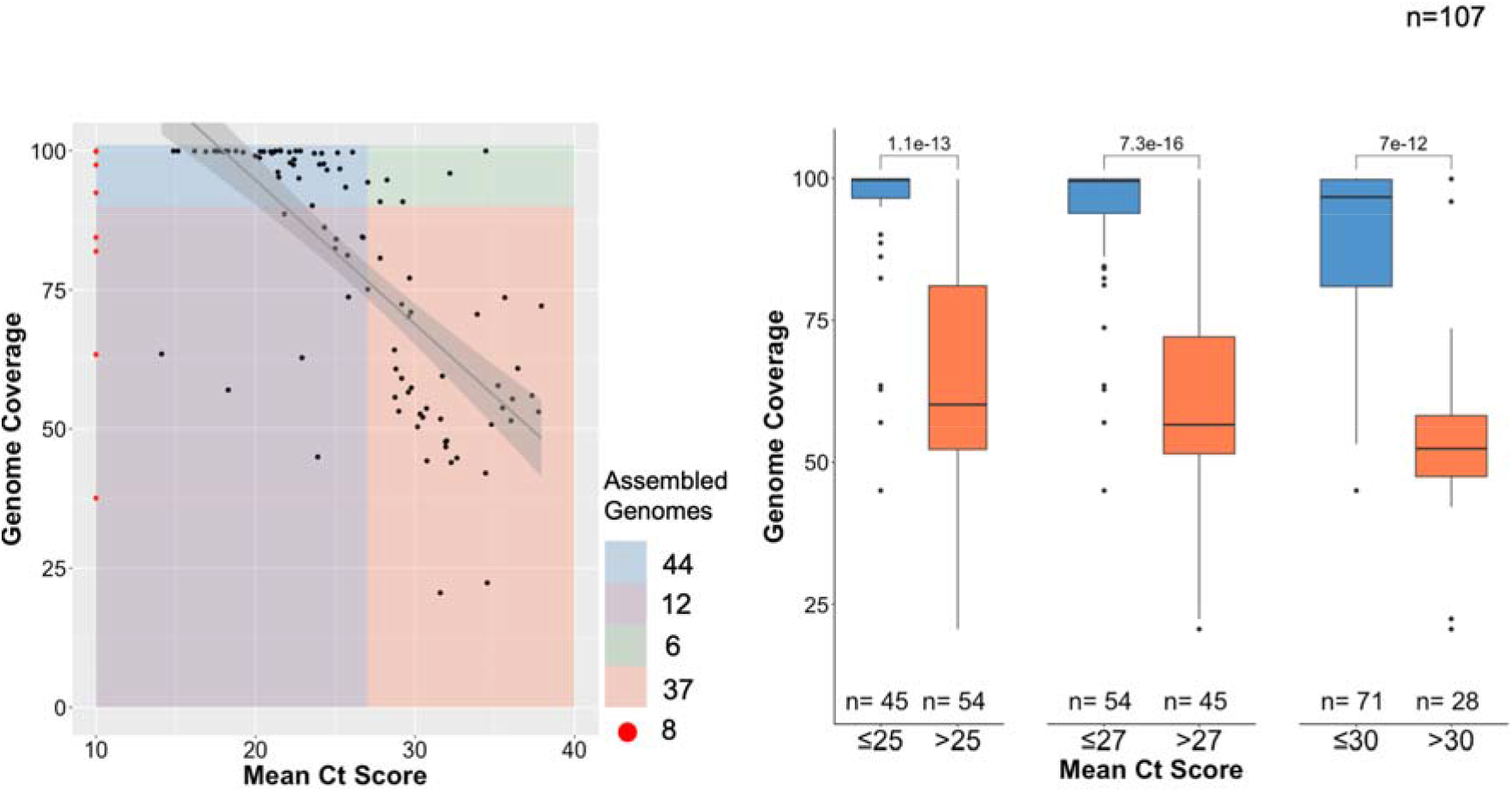
Association between Ct Score and genome length. A) Regression plot of mean Ct score of all unique samples against their genome lengths (% coverage against SARS-CoV-2 reference). Samples with missing Ct score information (n=8) are shown in red. We produced 44 assembled genomes of >90% from samples having Ct score <27 (blue), 6 genomes of >90% and Ct score >27 (green), 12 genomes <90% coverage and Ct score <27 (purple), and 37 genomes <90% coverage and Ct score >27 (orange). B) Box plot and statistical comparison of genome coverage obtained from samples grouped by 3 mean Ct score thresholds (25, 27, 30), showing statistically significant (t-tests) differences between lower and higher Ct score samples.

### Comparison of Library Preparation Methods

The three different library preparation methods, i.e. TruSeq DNA, NEBNext Ultra II and Nextera DNA Flex, produced similar results. In order to properly evaluate the performance of the library methods, we generated a subset of multiple sequencing for 12 samples, four of which were sequenced in triplicate and four in duplicate for each of the two new library preparation methods (Table 1). For example, ten of the twelve sequences produced with the different methods provided very similar genome coverage and mutations. In addition, using a Ct Score lower than 30, the TruSeq DNA and NEBNext Ultra II protocols produced 14 and 40 whole-genomes, which represented 66.7% and 45.3% of the genomes produced, respectively.

**Table 1.** Comparison of coverage and Ct scores between the different library preparation methods for repeat samples only.

| Sequence | Coverage (% of SARS-CoV-2 genome) |  |  | Ct Score |
| --- | --- | --- | --- | --- |
|  | TruSeq DNA Nano | NEBnext Ultra II | Nextera Flex |  |
| KRISP_0002 | 97.5 | 98.3 | 98.5 | 24 010 |
| KRISP_0004 | 99.5 | 98.3 | 98.1 | 24 143 |
| KRISP_0019 | 97.4 | <u>89.9</u> | 90.2 | NA |
| KRISP_0021 | <u>63.5</u> | 99.1 | <u>82.7</u> | 14 112 |
| KRISP_0024 | - | 95.2 | 94.6 | 21 452 |
| KRISP_0026 | - | 99.8 | 99.9 | 17 981 |
| KRISP_0028 | - | 96.1 | 98.1 | 21 399 |
| KRISP_0031 | - | 86.2 | 84.1 | 24 318 |
| KRISP_016 | 99.8 | - | 99.2 | NA |
| KRISP_017 | 99.9 | - | 99.9 | NA |
| KRISP_006 | 99.5 | - | 96.3 | 21 022 |
| KRISP_007 | 99.9 | - | 99.6 | 25 522 |
| KRISP_003 | 25.3 | 92.4 | - | Undetermined |
| KRISP_010 | 93.4 | 92.3 | - | 25 647 |
| KRISP_014 | 81.9 | 70.6 | - | NA |
| KRISP_013 | 63.4 | 73.0 | - | NA |

The comparison of cost and hands-on time initially showed that the NEBnext Ultra II provided the most cost-effective solution, as all components cost approximately $42 per genome and the hands-on time was six hours. The Nextera Flex is the easiest and quickest method, with a three hours hands-on time, but it cost approximately $60 per genome at full reaction usage. The hands-on time of TruSeq, the original library in the ARTIC protocol, was twelve hours and the cost for us was $52 per reaction. It is important to note, that prices in South Africa are higher than most regions in the world and are deeply affected by currency fluctuations. In addition, some kits require additional components that need to be purchased. The Illumina Nextera DNA Flex and the TruSeq Nano DNA Library Preparation kits contains all the reagents required to perform the library preparation. The NEBNext Ultra II DNA kit does not include some of the reagents required for library preparation, such as the end repair reagents, NEBNext Ultra II End Repair/dA-Tailing Module (New England BioLabs, Massachusetts, USA) and Ampure beads (Beckman Coulter, High Wycombe, UK). However, if one puts the cost of the personnel and the extra reagents needed for the NEBNext Ultra II, the Nextera Flex seems to be the most cost-effective and easy to use library preparation kit.

### Phylogenetic and lineage analysis for near full length genomes

We performed a very basic phylogenetic analysis of the near full length genomes as this analysis is beyond the scope of this paper. The 54 near full length genomes belong to lineages B(n=3), B.1. (n=50) and lineage B.2. (n=1) (Figure 4, Supplementary Table S2). All of the outbreak sequences clustered closely to each other in the lineage B.1. and they were 99.99% identical, with one or two mutations that differentiate themselves. The number of mutations (mean of 5 mutations from the original Wuhan reference sequences) of most of our sequences were in line with other public sequences sampled at the same time, furthermore, our outbreak sequences had an specific mutation, 16736 C to T (Supplementary Table S2), which caused a non-synonymous mutation at the ORFab1 gene position. More detailed phylogenetic analysis are described in our early genomic epidemiology report of SARS-CoV-2 infection KZN and our investigation of the hospital outbreak [6,13].

**Figure 4:**
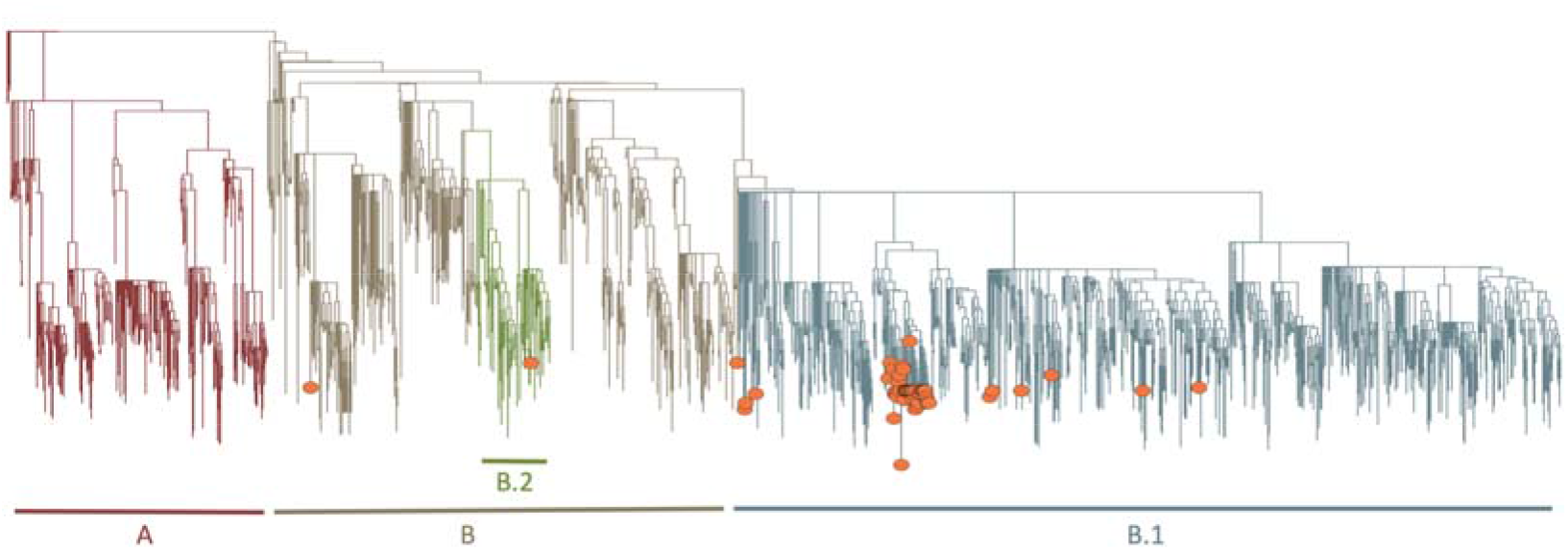
Phylogenetic tree. Showing a Maximum-Likelihood (ML) tree of our 54 genomes (orange circles) against publicly available SARS-CoV-2 genomes as reference. Our genomes fall mostly in the B.1 (n=50), B (n=3) or B.2 (n=1) lineages.

## Discussion

SARS-CoV-2 sequencing is a rapid and accurate method of analysing genetic variability and identifying transmission chains during outbreaks [15]. However, as outbreaks happen sporadically and cannot be predicted, it is not always possible to have all the resources required to perform the necessary tests in resource limited settings. As in many countries, there was a national lockdown in South Africa during the SARS-CoV-2 pandemic, with limited flights in and out of the country, thus making it difficult for suppliers to deliver reagents. In addition, reagents for SARS-CoV-2 were in great demand in the world making them more difficult to access.

All samples from the outbreak investigation had initially been collected and tested at different private and government laboratories. Results were required urgently for genomic sequencing and therefore tracing the exact qPCR results from the original labs to determine the cycling threshold (Ct) scores was impractical. Ideally, as shown in other reports of SARS-CoV-2 genomic epidemiology [8] samples should be tested on qPCR first. Those with a Ct below 30 have been shown to produce longer and higher quality genomes. A recent study of high-throughput sequencing in Iceland showed up to 90% success rate of sequencing SARS-CoV-2 near full length genomes for samples with Ct < 30. However, this was a prospective study in a very developed country. We found that in our setting and using samples that had been not optimally collected a Ct score of < 27 provided the best cut-off to produce near full length genomes.

In this study, we started using the ARTIC method [14] to amplify the SARS-CoV-2 using a tiling PCR approach. This method recommends the use the TruSeq DNA Nano kit for Illumina sequencing, which is a very labour intensive approach (up to 12h hands on time). As we ran out of reagents for TruSeq and needed to set up other methods quickly, we found a number of much better library sequencing kits that were either cheaper or were much less laborious. We evaluated and used the NEBnext Ultra II (New England BioLabs) and Nextera DNA Flex (Illumina). The NEBnext kit was used to generate the majority of the 108 genomes and 54 near full-length genomes. We also evaluated the Nextera Flex DNA library preparation kit, which saves up to 9h of hands on time, when compared with the original ARTIC protocol that uses TruSeq.

All three library preparation methods produced high quality genomes. There was no significant difference seen in coverage. The marginally lower coverage seen with the NEBnext Ultra II kit could be due to the unavailability of the recommended adapters and the substitution with the TruSeq adapters. Slightly lower coverage seen with Nextera Flex DNA could be due to the samples being subjected to freeze thaw cycles as the Nextera Flex DNA kit was used a month after samples were received and tested using the other two kits.

Further comparison of the kits looked at the cost and processing times for each of the methods used. The costs of the library preparation methods, including components not included, such as Ampure bead (Beckman Coulter) and End Repair/dA-Tailing Module (New England BioLabs) kits, varied with a difference of up to 30% (Supplementary Table S4). The NEBNext Ultra II DNA library method was found to be the cheaper option. Nextera DNA Flex was found to have the shortest processing time of less than three hours. While all three methods required end repair of amplicons prior to indexing, the Nextera Flex method encompassed the tagmentation step together with the ligation of adapters. There was no need to quantify and normalise individual libraries at the end of library preparation as normalization occurred during the tagmentation step.

Our study has many limitations. Firstly, we did not have time to prepare properly for the initial sequencing as our access to the first positive samples was during a large nosocomial outbreak investigation. Secondly, the quality of the samples was not homogeneous, as some samples arrived at our laboratories weeks after being sampled from the patients. Thirdly, reagents stockouts were common during the lockdown in South Africa and we had to innovate and adapt the protocols.

To summarise, despite the difficulties posed by the lockdown, we were able to complete the data generation and analysis of a large COVID-19 outbreak in South Africa in just a few weeks. We also evaluated the performance of three library preparation kits for their quality, cost, ease of use and time efficiency. In addition, we adapted a bioinformatics workflow to assemble SARS-CoV-2 genomes from raw sequence reads in near-real time. All of our protocols and raw data have been made publicly available and distributed to laboratories of the South African Network for Genomics Surveillance (SANGS) and the Africa Centre for Diseases Control (Africa CDC).

## Materials and Methods

### Sample collection and preparation

We obtained remnant samples collected using nasopharyngeal and oropharyngeal swabs from COVID-19 confirmed patients. These samples consisted of either the primary swab sample or extracted RNA. We heated inactivated swab samples in a water bath at 60°C for 30 minutes, prior to RNA extraction. We extracted RNA on an automated Chemagic 360 instrument using the CMG-1049 kit (Perkin Elmer, Hamburg, Germany), or by manual extraction using the QIAamp Viral RNA Mini Kit (QIAGEN, California, USA). The RNA was stored at −80 C prior to use.

### Real-time polymerase chain reaction

We retrieved RNA samples stored at −80°C and thawed them to room temperature prior to real-time PCR testing. We tested for three SARS-CoV-2 genes, i.e. ORF1ab, S protein and N protein, using the TaqMan 2019-nCoV assay kit v1 according to manufacturer’s instructions. In summary, we prepared a 20μl mastermix for each target gene (i.e. ORF1ab, S protein and N protein) containing; 11.25μl of nuclease free water, 6.25μl of TaqMan Fast Virus 1-Step Master Mix (4X), 1.25μl RNase P Assay (20X), and 1.25μl of the 2019-nCoV assay (20X), in respective tubes. We added 20μl of the mastermix to a 96-well plate and 5μl of RNA to the respective wells, and included a positive control (i.e. 1μl TaqMan 2019-nCoV Control Kit v1 and 4μl of nuclease free water) and no template control (i.e. 5μl nuclease free water) for each target gene.

We performed qPCR on a QuantStudio 7 Flex Real-Time PCR instrument (Life Technologies, Carlsbad, CA) using the following conditions; 50°C for five minutes, 95°C for 20 seconds, 40 cycles of 95°C for three seconds, and 60°C for 30 seconds. We analysed cycle thresholds (Ct) using auto-analysis settings with the threshold lines falling within the exponential phase of the fluorescence curves and above any background signal. To accept the results, we confirmed a Ct value for RNAse P (i.e. an endogenous internal amplification control) and or the target gene in each reaction, with undetermined Ct values in the no template control. We reported Ct values for each target gene.

### Tiling-based polymerase chain reaction

We performed cDNA synthesis using SuperScript IV reverse transcriptase (Life Technologies) and random hexamer primers, followed by gene specific multiplex PCR using the ARTIC protocol, as described previously [14]. In summary, we attempted SARS-CoV-2 whole genome amplification by multiplex PCR using primers designed on Primal Scheme (http://primal.zibraproject.org/) to generate 400 base pair (bp) amplicons with 70bp overlaps, covering the 30 kilobase SARS-CoV-2 genome. We purified PCR products in a 1:1 ratio using AmpureXP purification beads (Beckman Coulter, High Wycombe, UK), and quantified the purified amplicon using the Qubit dsDNA High Sensitivity assay kit on a Qubit 4.0 instrument (Life Technologies). We estimated amplicon fragment sizes on a LabChip GX Touch (Perkin Elmer, Hamburg, Germany) prior to library preparation.

### Library preparation and next generation sequencing

Depending on availability of reagents, we attempted library preparation using three different kits, namely TruSeq DNA Library Prep kit, NEBNEXT Ultra II DNA Library Prep Kit, and the Nextera DNA Flex Library Prep kit.

### TruSeq Nano DNA Library Preparation

We used the TruSeq Nano DNA library preparation kits (Illumina, San Diego, USA) to prepare uniquely indexed paired end libraries of genomic DNA according to the manufacturer’s instructions. In summary, we used 200ng of input DNA from purified amplicons for end-repair reaction, or 10μl of neat sample in cases of insufficient concentration. We performed adapter ligation using Illumina TruSeq Single DNA Indexes Set A and Set B (Illumina, San Diego, USA), and performed adaptor enrichment using (8) eight cycles of PCR. We quantified the libraries using the Qubit dsDNA High Sensitivity assay kit on a Qubit 4.0 instrument (Life Technologies, California, USA) and analysed the fragment sizes using a LabChip GX Touch (Perkin Elmer, Hamburg, Germany). We normalized each sample library to 4nM concentration, pooled the normalized libraries, and denatured them with 0.2N sodium acetate. We spiked 12 pM library with 1% PhiX control (PhiX Control v3) and sequenced on an Illumina MiSeq platform (Illumina, San Diego, USA) using a MiSeq Nano Reagent Kit v2 (500 cycles). We were limited from using this method further due to insufficient reagents available for the library preparation.

### NEBNext Ultra II DNA Library Preparation

We used the NEBNext Ultra II DNA Library Preparation kits according to the manufacturer’s instructions. In summary, we diluted purified tiling PCR amplicons to 5ng/μl and performed End Prep reactions on the amplicons using NEBNext Ultra II End Repair/dA-Tailing Module (New England BioLabs, Massachusetts, USA). Due to the national lockdown, we had a reagent stockout of the NEBNext Multiplex Oligos from Illumina. As a result, we performed adapter ligation using remaining Illumina TruSeq Single DNA Indexes Set A and Set B (Illumina, San Diego, USA), and performed adapter enrichment using the remaining PCR Primer Cocktail from the Illumina TruSeq Nano DNA Library Prep kit. We adapted PCR cycling conditions accordingly for the reagent substitution, and used eight cycles for the enrichment step. We performed size selection using 0.9x AmpureXP purification beads (Beckman Coulter, High Wycombe, UK), and quantified the libraries using the Qubit dsDNA High Sensitivity assay kit on a Qubit 4.0 instrument (Life Technologies, California, USA). We analysed the fragment sizes using a LabChip GX Touch (Perkin Elmer, Hamburg, Germany), with expected fragments between 300-600bp in size. We normalized each sample library to 4nM concentration, pooled the normalized libraries, and denatured them with 0.2N sodium acetate. We spiked in 1% PhiX control (PhiX Control v3) in a 12 pM library and sequenced on an Illumina MiSeq platform (Illumina, San Diego, CA, USA) using a MiSeq Nano Reagent Kit v2 (500 cycles).

### Nextera DNA Flex Library Preparation

We used the Nextera DNA Flex Library Prep kits (Illumina, San Diego, USA) according to the manufacturer’s instructions. We used undiluted tiling PCR amplicons. Briefly, we tagmented the DNA with bead-linked transposomes and stopped the tagmentation reaction with tagmentation stop buffer before proceeding to the post tagmentation cleanup using the tagmentation wash buffer. This step was followed by eight cycles of amplification of the tagmented DNA with enhanced PCR mix and index adapters. We used the Nextera DNA CD indexes (Illumina, San Diego, USA). We cleaned the libraries using 0.9x sample purification beads and eluted in 32μl resuspension buffer. Libraries were quantified using the Qubit dsDNA High Sensitivity assay kit on a Qubit 4.0 instrument (Life Technologies, California, USA). We analysed the fragment sizes using a LabChip GX Touch (Perkin Elmer, Hamburg, Germany), with expected fragments between 500-600bp in size. We normalized each sample library to four nM concentration, pooled the normalized libraries, and denatured them with 0.2N sodium acetate. We spiked the 12pM library with 1% PhiX control (PhiX Control v3) and sequenced on an Illumina MiSeq platform (Illumina, San Diego, USA) using a MiSeq Nano Reagent Kit v2 (500 cycles).

### Data analysis

Raw reads from Illumina sequencing were assembled using Genome Detective 1.126 (https://www.genomedetective.com/) and the coronavirus typing tool [16,17]. The initial assembly obtained from Genome Detective was polished by aligning mapped reads to the references and filtering out mutations with low genotype likelihoods using bcftools 1.7-2 mpileup method. All mutations were confirmed visually with bam files using Geneious software. For samples with repeat sequencing, forward and reverse reads from all sequencing runs were merged respectively and assembled as one. All of the sequences were deposited in GISAID (https://www.gisaid.org/). Lineage assignments were established using a dynamic lineage classification method proposed by Rambault et al., [18] via the Phylogenetic Assignment of named Global Outbreak LINeages (PANGOLIN) software suite (https://github.com/hCoV-2019/pangolin). 10,959 GISAID reference genomes (All authors acknowledged in Supplementary Table S6) and 54 KRISP sequences were aligned in Mafft v7·313 (FF-NS-2) followed by manual inspection and editing in the Geneious Prime software suite (Biomatters Ltd, New Zealand). We constructed a maximum likelihood (ML) tree topology in IQ-TREE (GTR+G+I, no support) [19,20]. The resulting phylogeny was viewed and annotated in FigTree and ggtree. All of the data produced has been deposited in the GISAID (consensus genomes) and at the fastq short reads deposited at the Short Read Archive (SRA) with accession: https://www.ncbi.nlm.nih.gov/nuccore/NC045512

## Supporting information

Acknowledgement to data used

