## Supplementary Tables for "Whole Genome Sequencing of SARS-CoV-2: Adapting Illumina Protocols for Quick and Accurate Outbreak Investigation During a Pandemic"

**Supplementary materia**

**Supplementary Figure S1**

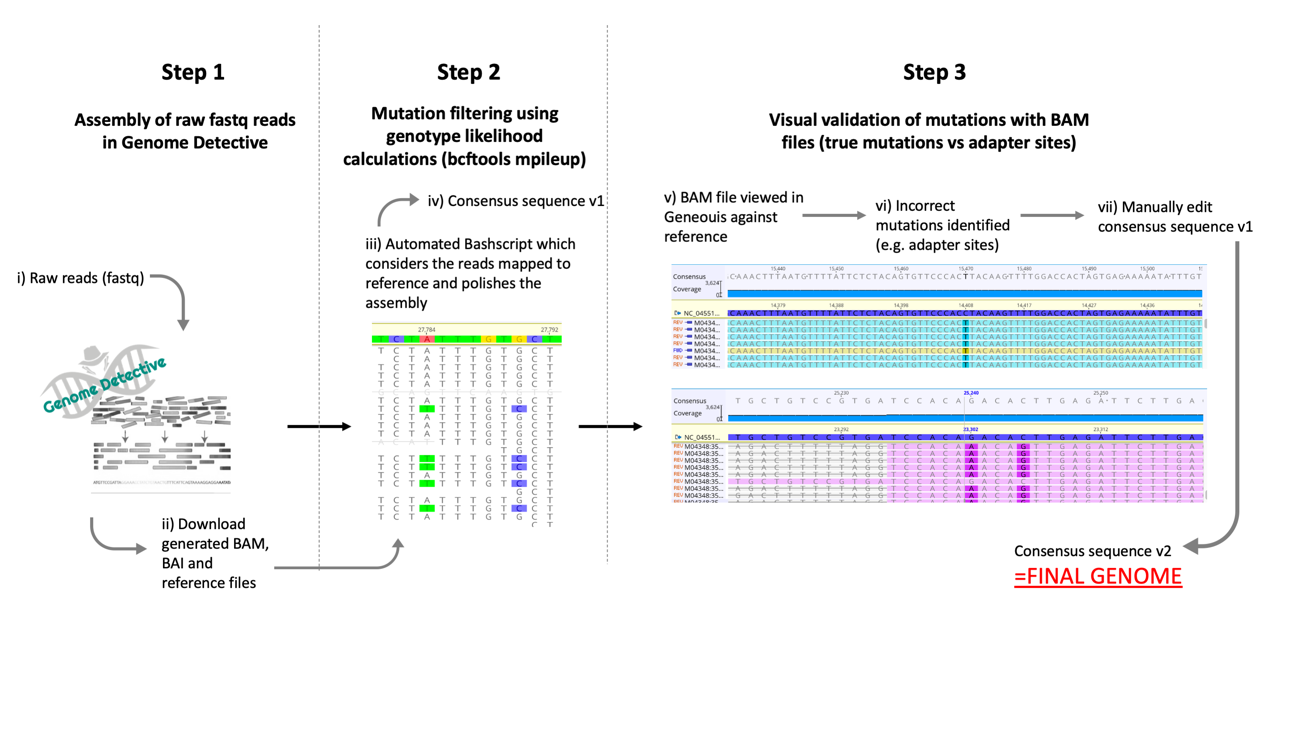

a)

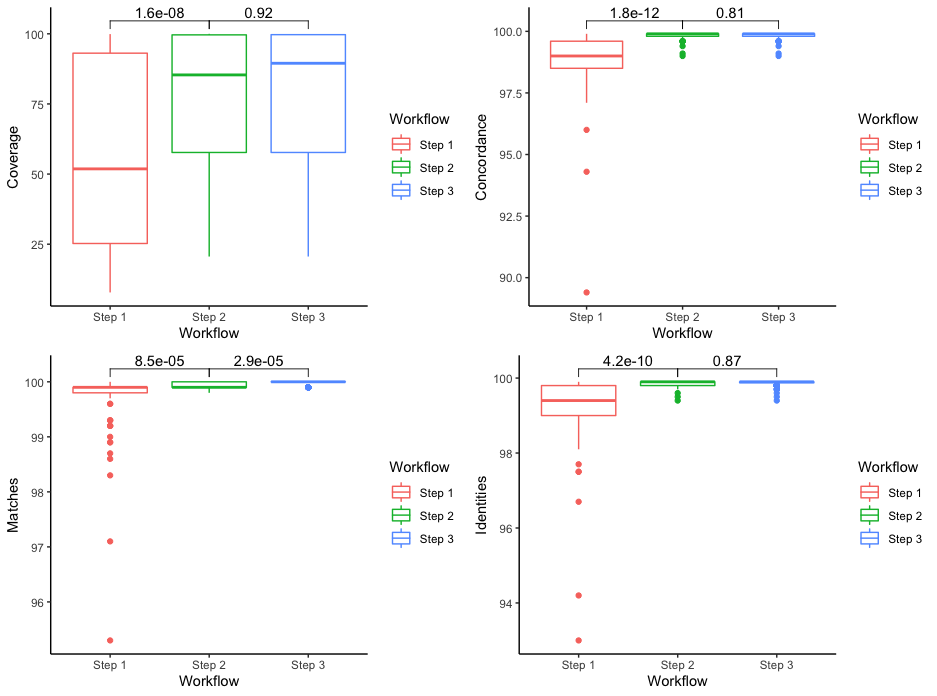

b)

**Supplementary Fig S1.** a) 3-step workflow for generation of high-quality genomes. Step 1: Raw reads were assembled using the web-based Genome Detective 1.126 (<https://www.genomedetective.com/>) platform and its coronavirus typing tool; Step 2: The initial assembly obtained from Genome Detective was polished by aligning mapped reads to the references and filtering out mutations with low genotype likelihoods using bcftools 1.7-2 mpileup method. This calculation determines the probability of a genotype at sites containing reads with various bases (e.g. the probability that position 27784 is A vs T in illustration above); Step 3: All mutations were validated visually with BAM files viewed in Geneious software to ensure that called mutations were true and not part of lingering adapter sites. b) Plot of genome coverage, concordance, matches, and identifies of our genomes against the reference at each stage of our 3-step workflow, showing significant improvement in all sequencing stats at step 2, compared to step 1 (t-test, p<0.05), and a significant increase in matches between step 2 and step 3 of the workflow.

**Supplementary Figure S2**

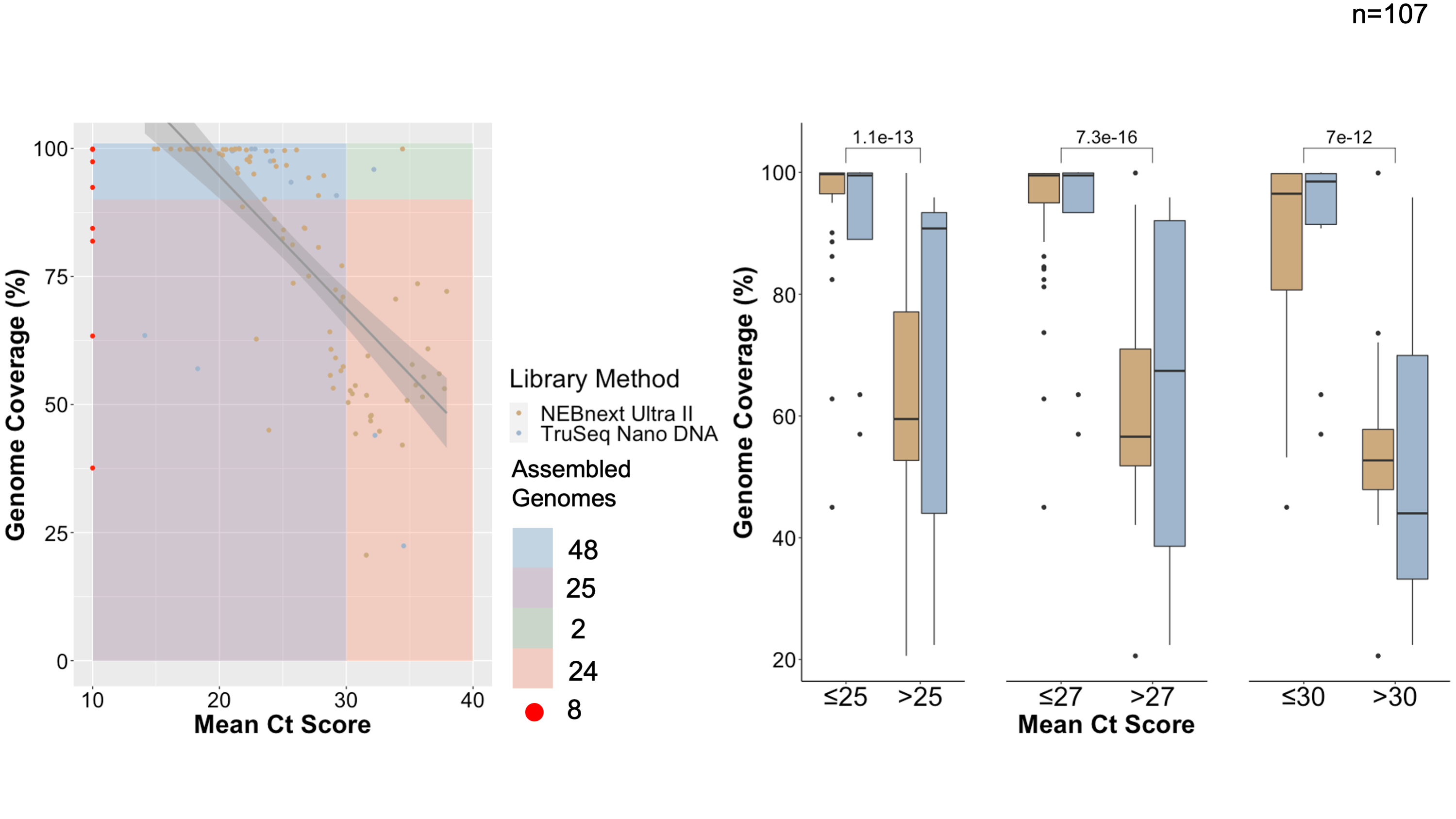

a)

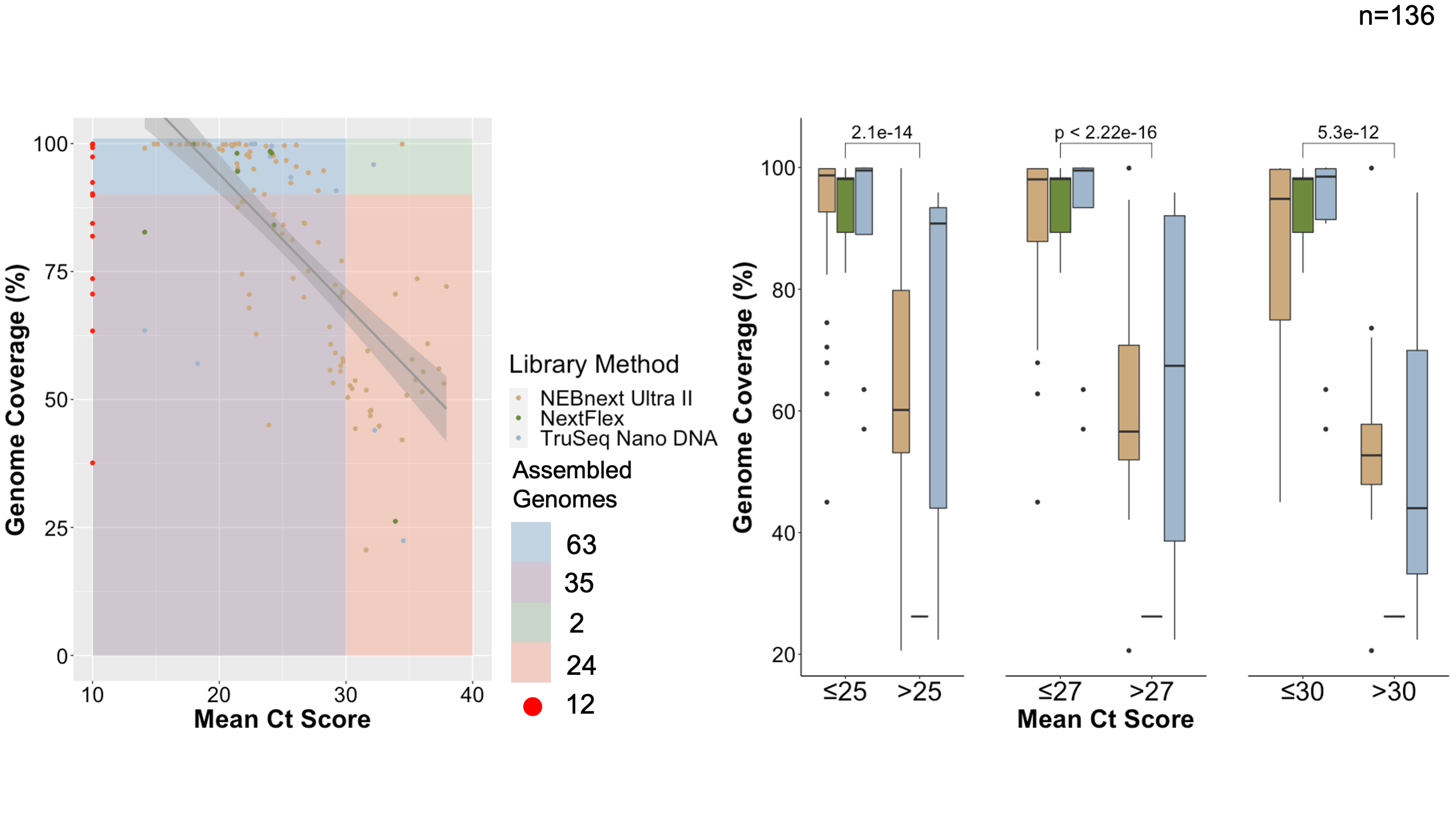

b)

**Supplementary Figure S2. Comparison of coverage obtained for the different library preparation methods against their mean ct scores.** Regression plot and box plots showing analysis of the effect of Ct score on final sequence coverage, color-coded by library method used, for 107 unique samples sequenced (a) and all 136 sequencing runs attempted, including repeats b). Color quadrants on regression plots show 4 categories of sequences, those with Ct score <30 and coverage >90% (blue), those with Ct score <30 and coverage <90% (purple), those with Ct score >30 and coverage >90% (green) and those with Ct score >30 and coverage <90% (orange). Sequences missing Ct score information are denoted in red

**Supplementary Table S1** – **Demographic and sequence information for 108 samples sequenced in this study.** Sequences selected for whole genome assembly in this study are highlighted in grey.

| **Sequence Name** | **Sampling date** | **Gender** | **Age** | **Ct_N_protein** | **Ct_ORF_1ab** | **Ct_S_protein** | **Average_Ct** | **Coverage** |
| --- | --- | --- | --- | --- | --- | --- | --- | --- |
| KRISP_0001 | 23/03/2020 | Female | 44 | 28373 | 30267 | 29050 | 29230 | 90.8 |
| KRISP_0002 | 23/03/2020 | Female | 40 | 23021 | 24695 | 24315 | 24010 | 97.5 |
| KRISP_0003 | 31/03/2020 |  |  |  |  |  |  | 92.4 |
| KRISP_0004 | missing | Female | 26 | 23144 | 25073 | 24211 | 24143 | 99.5 |
| KRISP_0005 | 31/03/2020 | Male | 30 | 31815 | 33722 | 31003 | 32180 | 95.9 |
| KRISP_0006 | 31/03/2020 | Male | 63 | 20389 | 23075 | 19601 | 21022 | 99.5 |
| KRISP_0007 | 01/04/2020 | Female | 44 | 21143 | 23455 | 22968 | 22522 | 99.9 |
| KRISP_0008 | 01/04/2020 |  |  | 32654 | 35338 | 35591 | 34528 | 22.4 |
| KRISP_0009 | 01/04/2020 |  |  | 31537 | 33062 | 32194 | 32264 | 44 |
| KRISP_0010 | 01/04/2020 | Female | 52 | 24582 | 26698 | 25661 | 25647 | 93.4 |
| KRISP_0011 | 01/04/2020 | Female | 40 | 20502 | 22443 | 21707 | 21551 | 100 |
| KRISP_0012 | 01/04/2020 | Female | 39 | 21524 | 23739 | 23175 | 22813 | 99.9 |
| KRISP_0013 | 25/03/2020 |  |  |  |  |  |  | 63.4 |
| KRISP_0014 | 02/04/2020 |  |  |  |  |  |  | 81.9 |
| KRISP_0015 | 02/04/2020 |  |  |  |  |  |  | 84.4 |
| KRISP_0016 | 28/03/2020 | Female | 57 |  |  |  |  | 99.8 |
| KRISP_0017 | missing | Male | 86 |  |  |  |  | 99.9 |
| KRISP_0018 | missing |  |  |  |  |  |  | 37.6 |
| KRISP_0019 | 02/04/2020 | Male | 80 |  |  |  |  | 97.4 |
| KRISP_0021 | 03/04/2020 | Male | 60 | 13101 | 14951 | 14285 | 14112 | 63.5 |
| KRISP_0022 | 03/04/2020 |  |  | 17662 | 18498 | 18700 | 18287 | 57 |
| KRISP_0023 | 02/04/2020 |  |  | 28274 | 29803 | 28849 | 28975 | 53.2 |
| KRISP_0024 | 02/04/2020 | Female | 26 | 20497 | 22174 | 21684 | 21452 | 95.2 |
| KRISP_0025 | 04/01/2020 |  |  | 29429 | 31032 | 29994 | 30152 | 50.4 |
| KRISP_0026 | 04/01/2020 | Female | 43 | 16604 | 19014 | 18326 | 17981 | 99.8 |
| KRISP_0027 | 02/04/2020 |  |  | 29634 | 31179 | 30134 | 30316 | 52.7 |
| KRISP_0028 | 02/04/2020 | Female | 49 | 20409 | 22242 | 21545 | 21399 | 96.1 |
| KRISP_0029 | 02/04/2020 | Female | 35 | 21814 | 23543 | 22798 | 22718 | 95 |
| KRISP_0030 | 02/04/2020 |  |  | 28427 | 30041 | 30271 | 29580 | 56.6 |
| KRISP_0031 | 02/04/2020 |  |  | 23409 | 25379 | 24166 | 24318 | 86.2 |
| KRISP_0032 | 02/04/2020 |  |  | 33394 | 34931 | 33361 | 33895 | 70.6 |
| KRISP_0033 | 01/04/2020 | Female | 38 | 19125 | 21195 | 20435 | 20252 | 98.7 |
| KRISP_0034 | 02/04/2020 |  |  | 31402 | 32975 | 31562 | 31980 | 47.9 |
| KRISP_0035 | 27/03/2020 | Female | 78* | 18188 | 20040 | 19436 | 19221 | 99.7 |
| KRISP_0036 | 27/03/2020 |  |  | 34558 | 35234 | 35816 | 35203 | 57.8 |
| KRISP_0037 | 27/03/2020 |  |  | 26617 | 28697 | 28123 | 27812 | 80.7 |
| KRISP_0038 | 27/03/2020 |  |  | 38106 | 34772 | 35439 | 36106 | 55.4 |
| KRISP_0039 | 27/03/2020 |  |  | 34980 | 36430 | 36631 | 36014 | 51.5 |
| KRISP_0040 | 27/03/2020 | Female | 86 | 33888 | 35417 | 34010 | 34438 | 99.9 |
| KRISP_0041 | 25/03/2020 | Female | 78* | 14142 | 15819 | 15451 | 15137 | 99.9 |
| KRISP_0042 | 01/04/2020 |  |  | 28101 | 29658 | 28624 | 28794 | 60.8 |
| KRISP_0043 | 01/04/2020 |  |  | 34215 | 35718 | 34449 | 34794 | 50.8 |
| KRISP_0044 | 01/04/2020 |  |  | 34618 | 35552 | 36279 | 35483 | 53.8 |
| KRISP_0045 | 27/03/2020 | Female | 39 | 15848 | 17568 | 17234 | 16883 | 99.8 |
| KRISP_0046 | 27/03/2020 |  |  | 24901 | 26619 | 25949 | 25823 | 73.7 |
| KRISP_0047 | 27/03/2020 |  |  | 39042 | 35619 |  | 37331 | 56 |
| KRISP_0048 | 27/03/2020 |  |  |  | 35452 | 37421 | 36437 | 60.9 |
| KRISP_0049 | 27/03/2020 |  |  | 37307 | 38169 |  | 37738 | 53.1 |
| KRISP_0050 | 28/03/2020 |  |  | 26165 | 27958 | 26977 | 27033 | 75.1 |
| KRISP_0051 | 28/03/2020 | Female | 55 | 22350 | 24498 | 23814 | 23554 | 90.1 |
| KRISP_0052 | 27/03/2020 |  |  | 28402 | 30011 | 29060 | 29158 | 59.1 |
| KRISP_0053 | 27/03/2020 |  |  | 37480 | 38362 |  | 37921 | 72.1 |
| KRISP_0055 | 31/03/2020 | Female | 82 | 17165 | 19038 | 18217 | 18140 | 99.9 |
| KRISP_0056 | 31/03/2020 | Male | 91* | 16736 | 18444 | 17915 | 17698 | 99.9 |
| KRISP_0057 | missing | Female | 35 | 25243 | 27180 | 25821 | 26081 | 99.7 |
| KRISP_0058 | missing | Male | 57 | 21015 | 22400 | 22007 | 21807 | 88.6 |
| KRISP_0059 | missing |  |  | 21557 | 22970 | 22724 | 22417 | 98.4 |
| KRISP_0060 | missing |  |  | 32217 | 32236 | 30666 | 31706 | 59.5 |
| KRISP_0061 | 03/04/2020 |  |  | 29595 | 30530 | 29087 | 29737 | 71 |
| KRISP_0062 | 03/04/2020 |  |  | 32486 | 32382 | 30853 | 31907 | 47.7 |
| KRISP_0063 | 04/04/2020 |  |  | 29366 | 30410 | 28913 | 29563 | 70.1 |
| KRISP_0064 | 04/04/2020 |  |  | 26915 | 27058 | 26079 | 26684 | 84.5 |
| KRISP_0065 | 04/04/2020 | Female | 32 | 21934 | 22676 | 21754 | 22121 | 99.7 |
| KRISP_0066 | 04/04/2020 | Male | 80 | 16331 | 16377 | 15806 | 16171 | 99.9 |
| KRISP_0067 | 03/04/2020 | Male | 70 | 15063 | 14927 | 14578 | 14856 | 99.9 |
| KRISP_0068 | 03/04/2020 |  |  | 32246 | 31951 | 30615 | 31604 | 51.8 |
| KRISP_0069 | 03/04/2020 |  |  | 23973 | 23294 | 21471 | 22913 | 62.8 |
| KRISP_0070 | 04/04/2020 |  |  | 29279 | 30466 | 29209 | 29651 | 77.1 |
| KRISP_0071 | 04/04/2020 |  |  | 30849 | 30847 | 29751 | 30482 | 52.1 |
| KRISP_0072 | 04/04/2020 |  |  | 32924 | 32665 | 32265 | 32618 | 44.8 |
| KRISP_0073 | 04/04/2020 |  |  | 26867 | 27173 | 26183 | 26741 | 84.4 |
| KRISP_0074 | 03/04/2020 | Female | 33 | 24439 | 24579 | 23817 | 24278 | 97.6 |
| KRISP_0075 | 04/04/2020 | Female | 40 | 20981 | 21292 | 20565 | 20946 | 99.8 |
| KRISP_0076 | 04/04/2020 |  |  | 35204 | 34829 | 33240 | 34424 | 42.1 |
| KRISP_0077 | 04/04/2020 |  |  | 30335 | 30131 | 28816 | 29761 | 57.4 |
| KRISP_0078 | 04/04/2020 |  |  | 29202 | 29151 | 27869 | 28741 | 55.7 |
| KRISP_0079 | 03/04/2020 |  |  | 28946 | 29215 | 27962 | 28708 | 64.2 |
| KRISP_0080 | missing |  |  | 21580 | 21409 | 20742 | 21244 | 99.9 |
| KRISP_0081 | missing | Female | 72 | 20177 | 22158 | 21258 | 21198 | 99.8 |
| KRISP_0082 | 06/04/2020 |  |  | 25154 | 26513 | 25631 | 25766 | 81.2 |
| KRISP_0083 | missing |  |  | 24453 | 25766 | 24947 | 25055 | 84.1 |
| KRISP_0084 | 05/04/2020 | Male | 55 | 19527 | 21411 | 20619 | 20519 | 99.8 |
| KRISP_0085 | 05/04/2020 |  |  | 30239 | 31513 | 30399 | 30717 | 53.7 |
| KRISP_0086 | 05/04/2020 |  |  | 30482 | 31471 | 30267 | 30740 | 44.3 |
| KRISP_0087 | missing |  |  | 23135 | 24641 | 23920 | 23899 | 45 |
| KRISP_0088 | missing | Female | 75 | 19201 | 16874 | 16605 | 17560 | 99.9 |
| KRISP_0089 | 07/04/2020 | Female | 64 | 21350 | 22969 | 22187 | 22169 | 97.8 |
| KRISP_0090 | 05/04/2020 | Female | 42 | 20970 | 23489 | 22680 | 22380 | 97.4 |
| KRISP_0091 | missing | Female | 87 | 19472 | 21043 | 20359 | 20291 | 99.8 |
| KRISP_0092 | 05/04/2020 |  |  | 34768 | 34934 | 37163 | 35622 | 73.6 |
| KRISP_0093 | missing |  |  | 24064 | 25874 | 25006 | 24981 | 82.4 |
| KRISP_0094 | missing |  |  | 29353 | 29557 | 28564 | 29158 | 72.4 |
| KRISP_0095 | missing |  |  | 31877 | 32820 | 31076 | 31924 | 46.8 |
| KRISP_0096 | missing |  |  | 20117 | 18624 | 21171 | 19971 | 99 |
| KRISP_0097 | missing |  |  | 17179 | 18878 | 18277 | 18111 | 99.8 |
| KRISP_0098 | missing |  |  | 31174 | 32626 | 30935 | 31578 | 20.6 |
| KRISP_0099 | missing |  |  | 26339 | 28024 | 26737 | 27033 | 94.3 |
| KRISP_0100 | missing |  |  | 26906 | 28896 | 27632 | 27811 | 90.8 |
| KRISP_0101 | missing |  |  | 21989 | 21303 | 21418 | 21570 | 99.9 |
| KRISP_0102 | 30/04/2020 | Female | 27 | 19173 | 18589 | 18564 | 18775 | 99.9 |
| KRISP_0103 | 29/04/2020 | Female | 23 | 25724 | 24785 | 24932 | 25147 | 99.6 |
| KRISP_0104 | 30/04/2020 | Male | 26 | 25604 | 25094 | 25146 | 25281 | 96.7 |
| KRISP_0105 | 30/04/2020 |  |  | 23891 | 23576 | 23591 | 23686 | 99.5 |
| KRISP_0106 | 30/04/2020 | Male | 61 | 18556 | 18109 | 18198 | 18288 | 99.9 |
| KRISP_0107 | 01/05/2020 | Male | 44 | 27697 | 28387 | 28626 | 28237 | 94.7 |
| KRISP_0108 | 04/05/2020 | Female | 45 | 23083 | 24898 | 25449 | 24477 | 96.5 |
| KRISP_0109 | 04/05/2020 | Male | 44 | 17692 | 17270 | 17332 | 17431 | 99.9 |

**Supplementary Table S2 – Change in sequencing statistics of 54 genomes generated in this study at 3 steps of the workflow. Final sequencing statistics for each genome highlighted in yellow.**

| **Sequence** | **Step** | **Begin** | **End** | **Coverage** | **Concordance** | **Matches** | **Identities** | **I/D** |
| --- | --- | --- | --- | --- | --- | --- | --- | --- |
| KRISP_001 | 1 | 27 | 29294 | 95 | 99,3 | 99,8 | 99,5 | 48/18 |
|  | 2 | 31 | 29867 | 99.60% | 99.90% | 29760 (99.9%) | 29754 (99.9%) | 0/12 |
|  | 3 | 31 | 29867 | 0.996 | 0.999 | 29772 (100%) | 29768 (99.9%) | 0/0 |
| KRISP_002 | 1 | 280 | 29488 | 82.30 | 99.70 | 24602 (99.9%) | 24574 (99.9%) | 2/3 |
|  | 2 | 46 | 29700 | 97.50% | 99.90% | 29145 (99.9%) | 29134 (99.9%) | 1/0 |
|  | 3 | 56 | 29700 | 97.40% | 99.90% | 29135 (100%) | 29129 (99.9%) | 0/0 |
| KRISP_003 | 1 | 321 | 29483 | 40.300 | 98.5000 | 99.9 | 99,2 | 3/0 |
|  | 2 | 31 | 29866 | 92.40% | 99.80% | 27632 (99.9%) | 27592 (99.8%) | 01-Mar |
|  | 3 | 31 | 29866 | 0.924 | 0.999 | 27635 (100%) | 27629 (99.9%) | 0/0 |
| KRISP_004 | 1 | 221 | 29857 | 83.00 | 99.70 | 24801 (99.9%) | 24758 (99.8%) | 0/6 |
|  | 2 | 46 | 29868 | 99.50% | 99.90% | 29756 (99.9%) | 29737 (99.9%) | 1/0 |
|  | 3 | 46 | 29868 | 99.50% | 99.90% | 29756 (100%) | 29742 (99.9%) | 0/0 |
| KRISP_005 | 1 | 30 | 29483 | 56.400 | 98.3000 | 98.3 | 97,5 | 6/274 |
|  | 2 | 27 | 29866 | 95.90% | 99.80% | 28690 (99.9%) | 28653 (99.9%) | 43831 |
|  | 3 | 27 | 29866 | 95.90% | 99.90% | 28691 (100%) | 28686 (99.9%) | 0/0 |
| KRISP_006 | 1 | 41 | 29866 | 97.80 | 99.70 | 29241 (99.9%) | 29203 (99.9%) | 1/1 |
|  | 2 | 13 | 29870 | 99.50% | 99.90% | 29742 (100%) | 29735 (99.9%) | 0/0 |
|  | 3 | 56 | 29867 | 99.30% | 99.90% | 29696 (100%) | 29692 (99.9%) | 0/0 |
| KRISP_007 | 1 | 7 | 29738 | 97.90 | 99.90 | 29287 (99.9%) | 29272 (99.9%) | 0/2 |
|  | 2 | 5 | 29875 | 99.90% | 99.90% | 29870 (99.9%) | 29861 (99.9%) | 0/1 |
|  | 3 | 56 | 29867 | 99.70% | 99.90% | 29812 (100%) | 29807 (99.9%) | 0/0 |
| KRISP_010 | 1 | 175 | 29489 | 71.90 | 99.40 | 21260 (98.9%) | 21206 (98.7%) | 2/230 |
|  | 2 | 171 | 29678 | 93.40% | 99.90% | 27943 (99.9%) | 27928 (99.9%) | 2/0 |
|  | 3 | 171 | 29678 | 93.40% | 99.90% | 27943 (100%) | 27933 (99.9%) | 0/0 |
| KRISP_011 | 1 | 104 | 29837 | 99.40 | 99.90 | 29734 (99.9%) | 29728 (99.9%) | 1/0 |
|  | 2 | 1 | 29903 | 100% | 99.90% | 29902 (99.9%) | 29894 (99.9%) | 1/1 |
|  | 3 | 56 | 29867 | 0.997 | 0.999 | 29812 (100%) | 29807 (99.9%) | 0/0 |
| KRISP_012 | 1 | 32 | 29857 | 95.00 | 99.70 | 28412 (99.9%) | 28369 (99.8%) | 0/8 |
|  | 2 | 4 | 29903 | 99.90% | 99.90% | 29872 (100%) | 29860 (99.9%) | 0/0 |
|  | 3 | 56 | 29867 | 0.996 | 0.999 | 29784 (100%) | 29773 (99.9%) | 0/0 |
| KRISP_016 | 1 | 112 | 29756 | 96.90% | 99.60% | 28963 (99.9%) | 28912 (99.8%) | 01-Feb |
|  | 2 | 1 | 29867 | 99.80% | 99.90% | 29835 (99.9%) | 29829 (99.9%) | 0/1 |
|  | 3 | 56 | 29867 | 0.997 | 0.999 | 29812 (100%) | 29807 (99.9%) | 0/0 |
| KRISP_017 | 1 | 104 | 29866 | 99.50% | 99.40% | 29760 (99.9%) | 29681 (99.6%) | 26/3 |
|  | 2 | 1 | 29892 | 99.90% | 99.90% | 29891 (99.9%) | 29886 (99.9%) | 0/1 |
|  | 3 | 56 | 29867 | 0.997 | 0.999 | 29812 (100%) | 29808 (99.9%) | 0/0 |
| KRISP_019 | 1 | 104 | 29756 | 83.00% | 99.70% | 24577 (99.0%) | 24539 (98.9%) | 1/239 |
|  | 2 | 46 | 29867 | 97.40% | 99.90% | 29112 (100%) | 29078 (99.9%) | 0/0 |
|  | 3 | 56 | 29867 | 0.997 | 0.999 | 29812 (100%) | 29808 (99.9%) | 0/0 |
| KRISP_021 | 1 | 47 | 29852 | 64.40% | 99.10% | 19256 (99.9%) | 19165 (99.5%) | 0/14 |
|  | 2 | 30 | 29867 | 99.70% | 99.90% | 29824 (100%) | 29816 (99.9%) | 0/0 |
|  | 3 | 30 | 29867 | 0.997 | 0.999 | 29824 (100%) | 29819 (99.9%) | 0/0 |
| KRISP_024 | 1 | 1043 | 29447 | 65.90% | 94.30% | 19015 (95.3%) | 18557 (93.0%) | 225/705 |
|  | 2 | 25 | 29866 | 97.10% | 99.90% | 29029 (99.9%) | 29023 (99.9%) | 1/0 |
|  | 3 | 25 | 29866 | 0.971 | 0.999 | 29029 (100%) | 29024 (99.9%) | 0/0 |
| KRISP_026 | 1 | 32 | 29827 | 96.00% | 99.80% | 28720 (99.9%) | 28692 (99.9%) | 43831 |
|  | 2 | 1 | 29870 | 99.80% | 99.90% | 29842 (100%) | 29834 (99.9%) | 0/0 |
|  | 3 | 1 | 29870 | 0.998 | 0.999 | 29842 (100%) | 29837 (99.9%) | 0/0 |
| KRISP_028 | 1 | 345 | 29831 | 77.40% | 99.50% | 23145 (99.9%) | 23098 (99.7%) | 43933 |
|  | 2 | 17 | 29868 | 96.10% | 99.90% | 28751 (99.9%) | 28743 (99.9%) | 2/0 |
|  | 3 | 17 | 29868 | 0.969 | 0.999 | 28975 (100%) | 28969 (99.9%) | 0/0 |
| KRISP_029 | 1 | 322 | 29459 | 71.10% | 99.20% | 21265 (99.9%) | 21189 (99.6%) | 10-Apr |
|  | 2 | 31 | 29866 | 95.00% | 99.90% | 28412 (99.9%) | 28404 (99.9%) | 0/1 |
|  | 3 | 31 | 29866 | 0.95 | 0.999 | 28413 (100%) | 28408 (99.9%) | 0/0 |
| KRISP_033 | 1 | 104 | 29789 | 92.40% | 99.50% | 27631 (99.9%) | 27561 (99.7%) | 03-Feb |
|  | 2 | 1 | 29871 | 98.70% | 99.90% | 29517 (99.9%) | 29506 (99.9%) | 0/1 |
|  | 3 | 1 | 29871 | 0.987 | 0.999 | 29518 (100%) | 29513 (99.9%) | 0/0 |
| KRISP_035 | 1 | 283 | 29866 | 92,1 | 99,7 | 99,9 | 99,8 | 0/14 |
|  | 2 | 9 | 29868 | 99.70% | 99.90% | 29810 (99.9%) | 29794 (99.9%) | 1/0 |
|  | 3 | 9 | 29868 | 0.999 | 0.999 | 29860 (100%) | 29856 (99.9%) | 0/0 |
| KRISP_040 | 1 | 47 | 29784 | 94,9 | 99,4 | 99,9 | 99,7 | 43958 |
|  | 2 | 13 | 29882 | 99.90% | 99.90% | 29870 (100%) | 29861 (99.9%) | 0/0 |
|  | 3 | 13 | 29882 | 0.999 | 0.999 | 29870 (100%) | 29866 (99.9%) | 0/0 |
| KRISP_041 | 1 | 31 | 29866 | 98,6 | 99,7 | 100 | 99,9 | 0/0 |
|  | 2 | 8 | 29870 | 99.90% | 99.90% | 29863 (100%) | 29859 (99.9%) | 0/0 |
|  | 3 | 8 | 29870 | 0.999 | 0.999 | 29863 (100%) | 29859 (99.9%) | 0/0 |
| KRISP_045 | 1 | 30 | 29750 | 99.20 | 99.9 | 99.9 | 99.9 | 0/8 |
|  | 2 | 30 | 29879 | 99.80% | 99.90% | 29850 (100%) | 29837 (99.9%) | 0/0 |
|  | 3 | 30 | 29879 | 99.80% | 99.90% | 29850 (100%) | 29838 (99.9%) | 0/0 |
| KRISP_051 | 1 | 322 | 29866 | 62.20 | 99.4 | 99.9 | 99.7 | 0/9 |
|  | 2 | 5 | 29868 | 90.10% | 99.90% | 26956 (100%) | 26936 (99.9%) | 0/0 |
|  | 3 | 5 | 29868 | 90.40% | 99.90% | 27041 (100%) | 27036 (99.9%) | 0/0 |
| KRISP_055 | 1 | 2 | 29866 | 99,9 | 99,8 | 99,9 | 99,9 | 0/1 |
|  | 2 | 1 | 29892 | 99.90% | 99.90% | 29892 (100%) | 29888 (99.9%) | 0/0 |
|  | 3 | 1 | 29892 | 0.999 | 0.999 | 29892 (100%) | 29888 (99.9%) | 0/0 |
| KRISP_056 | 1 | 4 | 29863 | 99,9 | 98,4 | 99,9 | 99,1 | 44123 |
|  | 2 | 1 | 29870 | 99.90% | 99.90% | 29870 (100%) | 29863 (99.9%) | 0/0 |
|  | 3 | 1 | 29870 | 0.999 | 0.999 | 29870 (100%) | 29864 (99.9%) | 0/0 |
| KRISP_057 | 1 | 821 | 29619 | 87,6 | 98,9 | 99,9 | 99,4 | 43892 |
|  | 2 | 31 | 29872 | 99.70% | 99.90% | 29809 (99.9%) | 29804 (99.9%) | 0/1 |
|  | 3 | 31 | 29872 | 0.997 | 0.999 | 29810 (100%) | 29806 (99.9%) | 0/0 |
| **Sequence** | **Step** | **Begin** | **End** | **Coverage** | **Concordance** | **Matches** | **Identities** | **I/D** |
| KRISP_058 | 1 | 322 | 29864 | 60 | 96 | 99,3 | 97,5 | 101/25 |
|  | 2 | 321 | 29866 | 92.90% | 99.80% | 27778 (99.9%) | 27749 (99.9%) | 01-Jan |
|  | 3 | 236 | 29866 | 0.932 | 0.999 | 27864 (100%) | 27859 (99.9%) | 0/0 |
| KRISP_059 | 1 | 32 | 29691 | 89,3 | 99,7 | 99,9 | 99,8 | 0 |
|  | 2 | 31 | 29866 | 98.40% | 99.90% | 29433 (99.9%) | 29421 (99.9%) | 1/0 |
|  | 3 | 31 | 29866 | 0.984 | 0.999 | 29433 (100%) | 29427 (99.9%) | 0/0 |
| KRISP_065 | 1 | 47 | 29712 | 95 | 99,3 | 99,9 | 99,6 | 44019 |
|  | 2 | 3 | 29866 | 99.70% | 99.90% | 29820 (100%) | 29805 (99.9%) | 0/0 |
|  | 3 | 3 | 29866 | 0.997 | 0.999 | 29820 (100%) | 29815 (99.9%) | 0/0 |
| KRISP_066 | 1 | 2 | 29866 | 98,1 | 99,7 | 99,9 | 99,8 | 43831 |
|  | 2 | 1 | 29875 | 99.90% | 99.90% | 29875 (100%) | 29866 (99.9%) | 0/0 |
|  | 3 | 1 | 29875 | 0.999 | 0.999 | 29875 (100%) | 29870 (99.9%) | 0/0 |
| KRISP_0067 | 1 | 104 | 29866 | 98,9 | 99,7 | 99,9 | 99,9 | 43834 |
|  | 2 | 1 | 29872 | 99.90% | 99.90% | 29872 (100%) | 29867 (99.9%) | 0/0 |
|  | 3 | 1 | 29872 | 0.999 | 0.999 | 29872 (100%) | 29868 (99.9%) | 0/0 |
| KRISP_074 | 1 | 37 | 29488 | 86,7 | 99,7 | 99,9 | 99,8 | 43900 |
|  | 2 | 31 | 29678 | 97.60% | 99.90% | 29171 (99.9%) | 29160 (99.9%) | 0/1 |
|  | 3 | 31 | 29678 | 0.976 | 0.999 | 29172 (100%) | 29167 (99.9%) | 0/0 |
| KRISP_075 | 1 | 39 | 29797 | 96,2 | 99,9 | 99,9 | 99,9 | 0/4 |
|  | 2 | 10 | 29869 | 99.80% | 99.90% | 29844 (100%) | 29838 (99.9%) | 0/0 |
|  | 3 | 10 | 29869 | 0.998 | 0.999 | 29844 (100%) | 29839 (99.9%) | 0/0 |
| KRISP_080 | 1 | 32 | 29799 | 96 | 99,8 | 99,9 | 99,9 | 43865 |
|  | 2 | 31 | 29903 | 99.90% | 99.90% | 29873 (100%) | 29861 (99.9%) | 0/0 |
|  | 3 | 31 | 29903 | 0.999 | 0.999 | 29873 (100%) | 29868 (99.9%) | 0/0 |
| KRISP_081 | 1 | 222 | 29836 | 95,2 | 99,6 | 99,9 | 99,8 | 43831 |
|  | 2 | 31 | 29868 | 99.80% | 99.90% | 29838 (99.9%) | 29826 (99.9%) | 1/0 |
|  | 3 | 31 | 29868 | 0.998 | 0.999 | 29838 (100%) | 29833 (99.9%) | 0/0 |
| KRISP_084 | 1 | 32 | 29691 | 95,6 | 99,7 | 99,8 | 99,7 | Aug-38 |
|  | 2 | 31 | 29866 | 99.80% | 99.90% | 29819 (99.9%) | 29808 (99.9%) | 0/17 |
|  | 3 | 31 | 29866 | 0.998 | 0.999 | 29836 (100%) | 29831 (99.9%) | 0/0 |
| KRISP_088 | 1 | 2 | 29866 | 99 | 99,9 | 99,9 | 99,9 | 0/4 |
|  | 2 | 1 | 29872 | 99.90% | 99.90% | 29872 (99.9%) | 29862 (99.9%) | 1/0 |
|  | 3 | 1 | 29872 | 0.999 | 0.999 | 29872 (100%) | 29867 (99.9%) | 0/0 |
| KRISP_089 | 1 | 47 | 29866 | 91 | 99,6 | 99,9 | 99,7 | 43872 |
|  | 2 | 46 | 29869 | 97.80% | 99.90% | 29246 (99.9%) | 29232 (99.9%) | 02-Jan |
|  | 3 | 46 | 29869 | 0.978 | 0.999 | 29247 (100%) | 29242 (99.9%) | 0/0 |
| KRISP_090 | 1 | 222 | 29836 | 85 | 99 | 99,9 | 43929 | 44062 |
|  | 2 | 30 | 29873 | 97.40% | 99.90% | 29111 (99.9%) | 29100 (99.9%) | 01-Oct |
|  | 3 | 30 | 29873 | 0.974 | 0.999 | 29121 (100%) | 29116 (99.9%) | 0/0 |
| KRISP_091 | 1 | 32 | 29690 | 95,4 | 99,9 | 99,9 | 99,9 | 0/6 |
|  | 2 | 31 | 29866 | 99.80% | 99.90% | 29836 (100%) | 29830 (99.9%) | 0/0 |
|  | 3 | 31 | 29866 | 0.998 | 0.999 | 29836 (100%) | 29831 (99.9%) | 0/0 |
| KRISP_096 | 1 | 17 | 29696 | 92,5 | 99,4 | 99,9 | 99,6 | 44181 |
|  | 2 | 17 | 29866 | 99.00% | 99.90% | 29616 (99.9%) | 29603 (99.9%) | 01-Jan |
|  | 3 | 17 | 29866 | 0.99 | 0.999 | 29617 (100%) | 29612 (99.9%) | 0/0 |
| KRISP_097 | 1 | 47 | 29425 | 96,9 | 99,8 | 99,9 | 99,9 | 0/1 |
|  | 2 | 1 | 29873 | 99.80% | 99.90% | 29854 (99.9%) | 29844 (99.9%) | 1/0 |
|  | 3 | 1 | 29873 | 0.998 | 0.999 | 29854 (100%) | 29849 (99.9%) | 0/0 |
| KRISP_099 | 1 | 47 | 28968 | 40,6 | 97,9 | 99,9 | 98,9 | 43925 |
|  | 2 | 34 | 29866 | 94.30% | 99.90% | 28191 (99.9%) | 28178 (99.9%) | 10-Mar |
|  | 3 | 34 | 29866 | 0.943 | 0.999 | 28194 (100%) | 28188 (99.9%) | 0/0 |
| KRISP_100 | 1 | 2 | 29061 | 47,8 | 98,9 | 99,9 | 99,4 | 43894 |
|  | 2 | 1 | 29866 | 90.80% | 99.80% | 27166 (100%) | 27131 (99.9%) | 0/0 |
|  | 3 | 1 | 29866 | 0.908 | 0.999 | 27166 (100%) | 27160 (99.9%) | 0/0 |
| KRISP_101 | 1 | 586 | 29756 | 97.000 | 99.9000 | 100 | 99,9 | 0/0 |
|  | 2 | 1 | 29869 | 0.999 | 0.9990 | 29869 (100%) | 29862 (99.9%) | 0/0 |
|  | 3 | 1 | 29869 | 0.999 | 0.999 | 29869 (100%) | 29864 (99.9%) | 0/0 |
| KRISP_102 | 1 | 32 | 29870 | 99.3 | 99.7 | 99.9 | 99,8 | 43831 |
|  | 2 | 1 | 29869 | 0.999 | 0.9990 | 29869 (100%) | 29860 (99.9%) | 0/0 |
|  | 3 | 1 | 29869 | 0.999 | 0.999 | 29869 (100%) | 29861 (99.9%) | 0/0 |
| KRISP_103 | 1 | 821 | 29493 | 85.400 | 98.9000 | 98.9 | 98,4 | 3/269 |
|  | 2 | 31 | 29866 | 0.996 | 0.9990 | 29798 (100%) | 29775 (99.9%) | 0/0 |
|  | 3 | 31 | 29866 | 0.996 | 0.999 | 29798 (100%) | 29790 (99.9%) | 0/0 |
| KRISP_104 | 1 | 321 | 29721 | 75.000 | 99.2000 | 99.9 | 99,6 | 43968 |
|  | 2 | 17 | 29869 | 0.967 | 0.9990 | 28917 (100%) | 28894 (99.9%) | 0/0 |
|  | 3 | 17 | 29869 | 0.967 | 0.9990 | 28917 (100%) | 28909 (99.9%) | 0/0 |
| KRISP_105 | 1 | 17 | 29493 | 84.400 | 99.3000 | 99.9 | 99,6 | 43932 |
|  | 2 | 21 | 29868 | 0.995 | 0.9990 | 29755 (99.9%) | 29743 (99.9%) | 0/6 |
|  | 3 | 21 | 29868 | 0.995 | 0.999 | 29761 (100%) | 29753 (99.9%) | 0/0 |
| KRISP_106 | 1 | 104 | 29866 | 99.500 | 99.7000 | 99.9 | 99,8 | 0/9 |
|  | 2 | 1 | 29871 | 0.999 | 0.9990 | 29871 (100%) | 29865 (99.9%) | 0/0 |
|  | 3 | 1 | 29871 | 0.999 | 0.9990 | 29871 (100%) | 29865 (99.9%) | 0/0 |
| KRISP_107 | 1 | 321 | 29864 | 37.900 | 98.4000 | 99.9 | 99,1 | 43933 |
|  | 2 | 31 | 29903 | 0.947 | 0.9980 | 28291 (99.8%) | 28267 (99.8%) | Mar-41 |
|  | 3 | 31 | 29903 | 0.949 | 0.9990 | 28364 (100%) | 28353 (99.9%) | 0/0 |
| KRISP_108 | 1 | 321 | 29691 | 54.100 | 98.8000 | 99.3 | 98,7 | 4/108 |
|  | 2 | 31 | 29868 | 0.965 | 0.9990 | 28848 (100%) | 28837 (99.9%) | 0/0 |
|  | 3 | 31 | 29868 | 0.965 | 0.9990 | 28848 (100%) | 28840 (99.9%) | 0/0 |
| KRISP_109 | 1 | 22 | 29872 | 99.800 | 99.7000 | 99.9 | 99,8 | 2/0 |
|  | 2 | 1 | 29885 | 0.999 | 0.9990 | 29885 (100%) | 29875 (99.9%) | 0/0 |
|  | 3 | 1 | 29885 | 0.999 | 0.9990 | 29885 (100%) | 29875 (99.9%) | 0/0 |

**Supplementary Table S3 – Mutations called, lineage assignment and GISAID accession ID for the SARS-CoV-2 genomes generated in this study**

| **Sequence ID** | **GISAID ID** | **Lineage Assignment** | **Number of mutations** | **Mutations** |
| --- | --- | --- | --- | --- |
| KRISP-001 |  | B | 4 | 241C>T, 3037C>T, 13536C>T, 14805C>T |
| KRISP-002 | EPI_ISL_421572 | B.2 | 6 | 4002C>T, 9223C>T, 11083G>T, 17247T>C, 26144G>T, 27927A>G |
| KRISP-003 | EPI_ISL_455629 | B.1 | 6 | 241C>T, 1877T>A, 3037C>T, 14408C>T,23404A>G, 24863C>A |
| KRISP-004 | EPI_ISL_436684 | B.1 | 9 | 241C>T, 3037C>T, 14408C>T, 18003T>C, 22675C>T, 23403A>G, 28881G>A, 28882G>A, 28883G>C |
| KRISP-005 | EPI_ISL_455630 | B.1 | 5 | 241C>T, 3037C>T, 14408C>T, 23403A>G, 26063G>T |
| KRISP-006 | EPI_ISL_421573 | B | 4 | 6312C>A, 11083G>T, 13730C>T, 28311C>T |
| KRISP-007 | EPI_ISL_421574 | B.1 | 5 | 241C>T, 3037C>T, 14408C>T, 16376C>T, 23403A>G |
| KRISP-010 | EPI_ISL_436685 | B.1 | 5 | 241C>T, 3037C>T, 14408C>T, 16376C>T, 23403A>G, |
| KRISP-011 | EPI_ISL_421575 | B.1 | 5 | 241C>T, 3037C>T, 14408C>T, 16376C>T, 23403A>G |
| KRISP-012 | EPI_ISL_421576 | B.1 | 11 | 241C>T, 3037C>T, 5220A>T, 14408C>T, 19170C>T, 19509G>A, 23403A>G, 25461T>C, 28881G>A, 28882G>A, 28883G>C |
| KRISP-016 |  | B.1 | 5 | 241C>T, 3037C>T, 14408C>T, 16376C>T, 23403A>G |
| KRISP-017 |  | B.1 | 4 | 241C>T, 3037C>T, 14408C>T, 23403A>G |
| KRISP-019 |  | B.1 | 4 | 241C>T, 3037C>T, 14408C>T, 23403A>G |
| KRISP-021 |  | B.1 | 5 | 241C>T, 2997C>T, 3037C>T, 14408C>T, 23403A>G |
| KRISP-024 |  | B.1 | 5 | 241C>T, 3037C>T, 14408C>T, 16376C>T, 23403A>G |
| KRISP-026 |  | B.1 | 5 | 241C>T, 3037C>T, 14408C>T, 16376C>T, 23403A>G |
| KRISP-028 |  | B.1 | 6 | 241C>T, 3037C>T, 14408C>T, 16376C>T, 19524C>T, 23403A>G |
| KRISP-029 |  | B.1 | 5 | 241C>T, 2388C>T, 3037C>T, 14408C>T, 23403A>G |
| KRISP-033 |  | B.1 | 5 | 241C>T, 3037C>T, 14408C>T, 16376C>T, 23403A>G |
| KRISP-035 |  | B.1 | 4 | 241C>T, 3037C>T, 14408C>T, 23403A>G |
| KRISP-040 |  | B.1 | 4 | 241C>T, 3037C>T, 14408C>T, 23403A>G |
| KRISP-041 |  | B.1 | 4 | 241C>T, 3037C>T, 14408C>T, 23403A>G |
| KRISP-045 | EPI_ISL_436686 | B.1 | 12 | 241C>T, 3037C>T, 4002C>T, 10097G>A, 11083G>T, 13536C>T, 14408C>T, 23403A>G, 23731C>T, 28881G>A, 28882G>A, 28883G>C |
| KRISP-051 | EPI_ISL_436687 | B.1 | 5 | 241C>T, 3037C>T, 14408C>T, 20161A>T, 23403A>G |
| KRISP-055 |  | B.1 | 4 | 241C>T, 3037C>T, 14408C>T, 23403A>G |
| KRISP-056 |  | B.1 | 6 | 241C>T, 856T>A, 1819A>C, 3037C>T, 14408C>T, 23403A>G |
| KRISP-057 |  | B.1 | 4 | 241C>T, 3037C>T, 14408C>T, 23403A>G |
| KRISP-058 |  | B.1 | 5 | 241C>T, 3037C>T, 14408C>T, 16376C>T, 23403A>G |
| KRISP-059 |  | B.1 | 6 | 241C>T, 1819A>C, 3037C>T, 14408C>T, 23403A>G, 23895C>T |
| KRISP-065 |  | B.1 | 5 | 241C>T, 3037C>T, 14408C>T, 16376C>T, 23403A>G |
| KRISP-066 |  | B.1 | 5 | 241C>T, 3037C>T, 14408C>T, 16376C>T, 23403A>G |
| KRISP-067 |  | B.1 | 4 | 241C>T, 3037C>T, 14408C>T, 23403A>G |
| KRISP-074 |  | B.1 | 5 | 241C>T, 3037C>T, 14408C>T, 16376C>T, 23403A>G |
| KRISP-075 |  | B.1 | 5 | 241C>T, 3037C>T, 14408C>T, 16376C>T, 23403A>G |
| KRISP-080 |  | B.1 | 5 | 241C>T, 3037C>T, 14408C>T, 16376C>T, 23403A>G |
| KRISP-081 |  | B.1 | 5 | 241C>T, 3037C>T, 14408C>T, 16376C>T, 23403A>G |
| KRISP-084 |  | B.1 | 5 | 241C>T, 3037C>T, 14408C>T, 16376C>T, 23403A>G |
| KRISP-088 |  | B.1 | 5 | 241C>T, 3037C>T, 14408C>T, 16376C>T, 23403A>G |
| KRISP-089 |  | B.1 | 5 | 241C>T, 3037C>T, 14408C>T, 16376C>T, 23403A>G |
| KRISP-090 |  | B.1 | 5 | 241C>T, 3037C>T, 14408C>T, 16376C>T, 23403A>G |
| KRISP-091 |  | B.1 | 5 | 241C>T, 3037C>T, 14408C>T, 16376C>T, 23403A>G |
| KRISP-096 |  | B.1 | 5 | 241C>T, 3037C>T, 14408C>T, 16376C>T, 23403A>G |
| KRISP-097 |  | B.1 | 5 | 241C>T, 3037C>T, 14408C>T, 16376C>T, 23403A>G |
| KRISP-099 |  | B | 6 | 241C>T, 273G>T, 343G>T, 8892C>T, 14408C>T, 23403A>G |
| KRISP-100 |  | B.1 | 6 | 241C>T, 440G>A, 3037C>T, 11083G>T, 14408C>T, 23403A>G |
| KRISP-101 | EPI_ISL_455631 | B.1 | 5 | 241C>T, 3037C>T, 8409T>C, 14408C>T, 23403A>G |
| KRISP-102 | EPI_ISL_455632 | B.1 | 8 | 241C>T, 3037C>T, 14408C>T, 23403A>G, 26004C>T, 28881G>A, 28882G>A, 28883G>C |
| KRISP-103 | EPI_ISL_455633 | B.1 | 8 | 241C>T, 3037C>T, 5672C>A, 10592A>G, 14408C>T, 16376C>T, 23403A>G, 26063G>T |
| KRISP-104 | EPI_ISL_455634 | B.1 | 8 | 241C>T, 3037C>T, 5672C>A, 10592A>G, 14408C>T, 16376C>T, 23403A>G, 26063G>T |
| KRISP-105 | EPI_ISL_455635 | B.1 | 8 | 241C>T, 3037C>T, 13766A>T, 14408C>T, 16376C>T, 18411T>C, 23403A>G, 24034C>T |
| KRISP-106 | EPI_ISL_455636 | B.1 | 6 | 241C>T, 3037C>T, 14408C>T, 16376C>T, 23403A>G, 24034C>T |
| KRISP-107 | EPI_ISL_455637 | B.1 | 11 | 241C>T, 3037C>T, 10950A>G, 14408C>T, 23191C>T, 23403A>G, 26884A>C, 26885C>A, 28881G>A, 28882G>A, 28883G>C |
| KRISP-108 | EPI_ISL_455638 | B.1 | 8 | 241C>T, 2364T>G, 3037C>T, 9086G>A, 14408C>T, 22800G>A, 23403A>G, 27171A>C |
| KRISP-109 | EPI_ISL_455639 | B.1 | 10 | 241C>T, 3037C>T, 8449A>C, 13115C>T, 14408C>T, 20234C>T, 23403A>G, 28881G>A, 28882G>A, 28883G>C |

**Supplementary Table S4 – Cost and logistics comparison of library methods tested in this study.**

|  | **TruSeq** | **NEBnext Ultra II** | **Nextera Flex** |
| --- | --- | --- | --- |
| Cost* | $ 52 | $ 42 | $ 60 |
| Time | 12 hours | 6 hours | 3 hours |
| Availability in Country** | Out of stock | Available | Out of stock |
| Lead time of Stock since order | 6 weeks | 2 days | 7 days |
| Local Supplier | Whitehead Scientific / Separations | Inqaba Biotec | Separations |

*Cost based on ZAR to USD exchange rate being R17.65 on the 25 May 2020 ([www.xe.com](http://www.xe.com)).

**Availability was reported at the time that the work was being performed in March 2020.

***At the time of purchasing the TruSeq DNA library preparation kit, Whitehead Scientific was the sole supplier of Illumina reagents in South Africa. During the write up of this manuscript, Separations had taken over being a sole supplier of Illumina reagents from April 2020. The Nextera Flex was obtained from Separations in April 2020.

**Supplementary Table S5 – PCR Primer Table (downloaded from ARTIC V3 protocol)**

| **Name** | **Pool** | **Sequence** | **Length** | **%GC** | **Tm (use 65)** |
| --- | --- | --- | --- | --- | --- |
| nCoV-2019_1_LEFT | nCoV-2019_1 | ACCAACCAACTTTCGATCTCTTGT | 24 | 41.67 | 60.69 |
| nCoV-2019_1_RIGHT | nCoV-2019_1 | CATCTTTAAGATGTTGACGTGCCTC | 25 | 44.00 | 60.45 |
| nCoV-2019_2_LEFT | nCoV-2019_2 | CTGTTTTACAGGTTCGCGACGT | 22 | 50.00 | 61.67 |
| nCoV-2019_2_RIGHT | nCoV-2019_2 | TAAGGATCAGTGCCAAGCTCGT | 22 | 50.00 | 61.74 |
| nCoV-2019_3_LEFT | nCoV-2019_1 | CGGTAATAAAGGAGCTGGTGGC | 22 | 54.55 | 61.32 |
| nCoV-2019_3_RIGHT | nCoV-2019_1 | AAGGTGTCTGCAATTCATAGCTCT | 24 | 41.67 | 60.32 |
| nCoV-2019_4_LEFT | nCoV-2019_2 | GGTGTATACTGCTGCCGTGAAC | 22 | 54.55 | 61.56 |
| nCoV-2019_4_RIGHT | nCoV-2019_2 | CACAAGTAGTGGCACCTTCTTTAGT | 25 | 44.00 | 60.97 |
| nCoV-2019_5_LEFT | nCoV-2019_1 | TGGTGAAACTTCATGGCAGACG | 22 | 50.00 | 61.39 |
| nCoV-2019_5_RIGHT | nCoV-2019_1 | ATTGATGTTGACTTTCTCTTTTTGGAGT | 28 | 32.14 | 60.17 |
| nCoV-2019_6_LEFT | nCoV-2019_2 | GGTGTTGTTGGAGAAGGTTCCG | 22 | 54.55 | 61.64 |
| nCoV-2019_6_RIGHT | nCoV-2019_2 | TAGCGGCCTTCTGTAAAACACG | 22 | 50.00 | 61.18 |
| nCoV-2019_7_LEFT | nCoV-2019_1 | ATCAGAGGCTGCTCGTGTTGTA | 22 | 50.00 | 61.73 |
| nCoV-2019_7_RIGHT | nCoV-2019_1 | TGCACAGGTGACAATTTGTCCA | 22 | 45.45 | 60.95 |
| nCoV-2019_8_LEFT | nCoV-2019_2 | AGAGTTTCTTAGAGACGGTTGGGA | 24 | 45.83 | 61.00 |
| nCoV-2019_8_RIGHT | nCoV-2019_2 | GCTTCAACAGCTTCACTAGTAGGT | 24 | 45.83 | 60.56 |
| nCoV-2019_9_LEFT | nCoV-2019_1 | TCCCACAGAAGTGTTAACAGAGGA | 24 | 45.83 | 61.18 |
| nCoV-2019_9_RIGHT | nCoV-2019_1 | ATGACAGCATCTGCCACAACAC | 22 | 50.00 | 61.71 |
| nCoV-2019_10_LEFT | nCoV-2019_2 | TGAGAAGTGCTCTGCCTATACAGT | 24 | 45.83 | 61.12 |
| nCoV-2019_10_RIGHT | nCoV-2019_2 | TCATCTAACCAATCTTCTTCTTGCTCT | 27 | 37.04 | 60.31 |
| nCoV-2019_11_LEFT | nCoV-2019_1 | GGAATTTGGTGCCACTTCTGCT | 22 | 50.00 | 61.66 |
| nCoV-2019_11_RIGHT | nCoV-2019_1 | TCATCAGATTCAACTTGCATGGCA | 24 | 41.67 | 61.35 |
| nCoV-2019_12_LEFT | nCoV-2019_2 | AAACATGGAGGAGGTGTTGCAG | 22 | 50.00 | 61.08 |
| nCoV-2019_12_RIGHT | nCoV-2019_2 | TTCACTCTTCATTTCCAAAAAGCTTGA | 27 | 33.33 | 60.36 |
| nCoV-2019_13_LEFT | nCoV-2019_1 | TCGCACAAATGTCTACTTAGCTGT | 24 | 41.67 | 60.56 |
| nCoV-2019_13_RIGHT | nCoV-2019_1 | ACCACAGCAGTTAAAACACCCT | 22 | 45.45 | 60.36 |
| nCoV-2019_14_LEFT | nCoV-2019_2 | CATCCAGATTCTGCCACTCTTGT | 23 | 47.83 | 60.62 |
| nCoV-2019_14_RIGHT | nCoV-2019_2 | AGTTTCCACACAGACAGGCATT | 22 | 45.45 | 60.42 |
| nCoV-2019_15_LEFT | nCoV-2019_1 | ACAGTGCTTAAAAAGTGTAAAAGTGCC | 27 | 37.04 | 61.32 |
| nCoV-2019_15_RIGHT | nCoV-2019_1 | AACAGAAACTGTAGCTGGCACT | 22 | 45.45 | 60.16 |
| nCoV-2019_16_LEFT | nCoV-2019_2 | AATTTGGAAGAAGCTGCTCGGT | 22 | 45.45 | 60.82 |
| nCoV-2019_16_RIGHT | nCoV-2019_2 | CACAACTTGCGTGTGGAGGTTA | 22 | 50.00 | 61.32 |
| nCoV-2019_17_LEFT | nCoV-2019_1 | CTTCTTTCTTTGAGAGAAGTGAGGACT | 27 | 40.74 | 60.69 |
| nCoV-2019_17_RIGHT | nCoV-2019_1 | TTTGTTGGAGTGTTAACAATGCAGT | 25 | 36.00 | 60.11 |
| nCoV-2019_18_LEFT | nCoV-2019_2 | TGGAAATACCCACAAGTTAATGGTTTAAC | 29 | 34.48 | 60.69 |
| nCoV-2019_18_RIGHT | nCoV-2019_2 | AGCTTGTTTACCACACGTACAAGG | 24 | 45.83 | 61.51 |
| nCoV-2019_19_LEFT | nCoV-2019_1 | GCTGTTATGTACATGGGCACACT | 23 | 47.83 | 61.18 |
| nCoV-2019_19_RIGHT | nCoV-2019_1 | TGTCCAACTTAGGGTCAATTTCTGT | 25 | 40.00 | 60.40 |
| nCoV-2019_20_LEFT | nCoV-2019_2 | ACAAAGAAAACAGTTACACAACAACCA | 27 | 33.33 | 60.68 |
| nCoV-2019_20_RIGHT | nCoV-2019_2 | ACGTGGCTTTATTAGTTGCATTGTT | 25 | 36.00 | 60.28 |
| nCoV-2019_21_LEFT | nCoV-2019_1 | TGGCTATTGATTATAAACACTACACACCC | 29 | 37.93 | 61.49 |
| nCoV-2019_21_RIGHT | nCoV-2019_1 | TAGATCTGTGTGGCCAACCTCT | 22 | 50.00 | 60.83 |
| nCoV-2019_22_LEFT | nCoV-2019_2 | ACTACCGAAGTTGTAGGAGACATTATACT | 29 | 37.93 | 61.25 |
| nCoV-2019_22_RIGHT | nCoV-2019_2 | ACAGTATTCTTTGCTATAGTAGTCGGC | 27 | 40.74 | 60.73 |
| nCoV-2019_23_LEFT | nCoV-2019_1 | ACAACTACTAACATAGTTACACGGTGT | 27 | 37.04 | 60.26 |
| nCoV-2019_23_RIGHT | nCoV-2019_1 | ACCAGTACAGTAGGTTGCAATAGTG | 25 | 44.00 | 60.57 |
| nCoV-2019_24_LEFT | nCoV-2019_2 | AGGCATGCCTTCTTACTGTACTG | 23 | 47.83 | 60.37 |
| nCoV-2019_24_RIGHT | nCoV-2019_2 | ACATTCTAACCATAGCTGAAATCGGG | 26 | 42.31 | 61.19 |
| nCoV-2019_25_LEFT | nCoV-2019_1 | GCAATTGTTTTTCAGCTATTTTGCAGT | 27 | 33.33 | 60.73 |
| nCoV-2019_25_RIGHT | nCoV-2019_1 | ACTGTAGTGACAAGTCTCTCGCA | 23 | 47.83 | 61.30 |
| nCoV-2019_26_LEFT | nCoV-2019_2 | TTGTGATACATTCTGTGCTGGTAGT | 25 | 40.00 | 60.28 |
| nCoV-2019_26_RIGHT | nCoV-2019_2 | TCCGCACTATCACCAACATCAG | 22 | 50.00 | 60.42 |
| nCoV-2019_27_LEFT | nCoV-2019_1 | ACTACAGTCAGCTTATGTGTCAACC | 25 | 44.00 | 60.80 |
| nCoV-2019_27_RIGHT | nCoV-2019_1 | AATACAAGCACCAAGGTCACGG | 22 | 50.00 | 61.13 |
| nCoV-2019_28_LEFT | nCoV-2019_2 | ACATAGAAGTTACTGGCGATAGTTGT | 26 | 38.46 | 60.13 |
| nCoV-2019_28_RIGHT | nCoV-2019_2 | TGTTTAGACATGACATGAACAGGTGT | 26 | 38.46 | 60.91 |
| nCoV-2019_29_LEFT | nCoV-2019_1 | ACTTGTGTTCCTTTTTGTTGCTGC | 24 | 41.67 | 61.39 |
| nCoV-2019_29_RIGHT | nCoV-2019_1 | AGTGTACTCTATAAGTTTTGATGGTGTGT | 29 | 34.48 | 60.69 |
| nCoV-2019_30_LEFT | nCoV-2019_2 | GCACAACTAATGGTGACTTTTTGCA | 25 | 40.00 | 61.19 |
| nCoV-2019_30_RIGHT | nCoV-2019_2 | ACCACTAGTAGATACACAAACACCAG | 26 | 42.31 | 60.30 |
| nCoV-2019_31_LEFT | nCoV-2019_1 | TTCTGAGTACTGTAGGCACGGC | 22 | 54.55 | 62.03 |
| nCoV-2019_31_RIGHT | nCoV-2019_1 | ACAGAATAAACACCAGGTAAGAATGAGT | 28 | 35.71 | 60.69 |
| nCoV-2019_32_LEFT | nCoV-2019_2 | TGGTGAATACAGTCATGTAGTTGCC | 25 | 44.00 | 61.09 |
| nCoV-2019_32_RIGHT | nCoV-2019_2 | AGCACATCACTACGCAACTTTAGA | 24 | 41.67 | 60.56 |
| nCoV-2019_33_LEFT | nCoV-2019_1 | ACTTTTGAAGAAGCTGCGCTGT | 22 | 45.45 | 61.58 |
| nCoV-2019_33_RIGHT | nCoV-2019_1 | TGGACAGTAAACTACGTCATCAAGC | 25 | 44.00 | 61.08 |
| nCoV-2019_34_LEFT | nCoV-2019_2 | TCCCATCTGGTAAAGTTGAGGGT | 23 | 47.83 | 61.02 |
| nCoV-2019_34_RIGHT | nCoV-2019_2 | AGTGAAATTGGGCCTCATAGCA | 22 | 45.45 | 60.03 |
| nCoV-2019_35_LEFT | nCoV-2019_1 | TGTTCGCATTCAACCAGGACAG | 22 | 50.00 | 61.39 |
| nCoV-2019_35_RIGHT | nCoV-2019_1 | ACTTCATAGCCACAAGGTTAAAGTCA | 26 | 38.46 | 60.69 |
| nCoV-2019_36_LEFT | nCoV-2019_2 | TTAGCTTGGTTGTACGCTGCTG | 22 | 50.00 | 61.44 |
| nCoV-2019_36_RIGHT | nCoV-2019_2 | GAACAAAGACCATTGAGTACTCTGGA | 26 | 42.31 | 60.74 |
| nCoV-2019_37_LEFT | nCoV-2019_1 | ACACACCACTGGTTGTTACTCAC | 23 | 47.83 | 60.93 |
| nCoV-2019_37_RIGHT | nCoV-2019_1 | GTCCACACTCTCCTAGCACCAT | 22 | 54.55 | 61.48 |
| nCoV-2019_38_LEFT | nCoV-2019_2 | ACTGTGTTATGTATGCATCAGCTGT | 25 | 40.00 | 60.86 |
| nCoV-2019_38_RIGHT | nCoV-2019_2 | CACCAAGAGTCAGTCTAAAGTAGCG | 25 | 48.00 | 61.13 |
| nCoV-2019_39_LEFT | nCoV-2019_1 | AGTATTGCCCTATTTTCTTCATAACTGGT | 29 | 34.48 | 61.00 |
| nCoV-2019_39_RIGHT | nCoV-2019_1 | TGTAACTGGACACATTGAGCCC | 22 | 50.00 | 60.55 |
| nCoV-2019_40_LEFT | nCoV-2019_2 | TGCACATCAGTAGTCTTACTCTCAGT | 26 | 42.31 | 61.25 |
| nCoV-2019_40_RIGHT | nCoV-2019_2 | CATGGCTGCATCACGGTCAAAT | 22 | 50.00 | 62.09 |
| nCoV-2019_41_LEFT | nCoV-2019_1 | GTTCCCTTCCATCATATGCAGCT | 23 | 47.83 | 60.75 |
| nCoV-2019_41_RIGHT | nCoV-2019_1 | TGGTATGACAACCATTAGTTTGGCT | 25 | 40.00 | 60.75 |
| nCoV-2019_42_LEFT | nCoV-2019_2 | TGCAAGAGATGGTTGTGTTCCC | 22 | 50.00 | 61.08 |
| nCoV-2019_42_RIGHT | nCoV-2019_2 | CCTACCTCCCTTTGTTGTGTTGT | 23 | 47.83 | 60.69 |
| nCoV-2019_43_LEFT | nCoV-2019_1 | TACGACAGATGTCTTGTGCTGC | 22 | 50.00 | 60.93 |
| nCoV-2019_43_RIGHT | nCoV-2019_1 | AGCAGCATCTACAGCAAAAGCA | 22 | 45.45 | 61.14 |
| nCoV-2019_44_LEFT | nCoV-2019_2 | TGCCACAGTACGTCTACAAGCT | 22 | 50.00 | 61.66 |
| nCoV-2019_44_RIGHT | nCoV-2019_2 | AACCTTTCCACATACCGCAGAC | 22 | 50.00 | 60.87 |
| nCoV-2019_45_LEFT | nCoV-2019_1 | TACCTACAACTTGTGCTAATGACCC | 25 | 44.00 | 60.57 |
| nCoV-2019_45_RIGHT | nCoV-2019_1 | AAATTGTTTCTTCATGTTGGTAGTTAGAGA | 30 | 30.00 | 60.01 |
| nCoV-2019_46_LEFT | nCoV-2019_2 | TGTCGCTTCCAAGAAAAGGACG | 22 | 50.00 | 61.38 |
| nCoV-2019_46_RIGHT | nCoV-2019_2 | CACGTTCACCTAAGTTGGCGTA | 22 | 50.00 | 60.86 |
| nCoV-2019_47_LEFT | nCoV-2019_1 | AGGACTGGTATGATTTTGTAGAAAACCC | 28 | 39.29 | 61.42 |
| nCoV-2019_47_RIGHT | nCoV-2019_1 | AATAACGGTCAAAGAGTTTTAACCTCTC | 28 | 35.71 | 60.06 |
| nCoV-2019_48_LEFT | nCoV-2019_2 | TGTTGACACTGACTTAACAAAGCCT | 25 | 40.00 | 61.09 |
| nCoV-2019_48_RIGHT | nCoV-2019_2 | TAGATTACCAGAAGCAGCGTGC | 22 | 50.00 | 60.74 |
| nCoV-2019_49_LEFT | nCoV-2019_1 | AGGAATTACTTGTGTATGCTGCTGA | 25 | 40.00 | 60.57 |
| nCoV-2019_49_RIGHT | nCoV-2019_1 | TGACGATGACTTGGTTAGCATTAATACA | 28 | 35.71 | 61.05 |
| nCoV-2019_50_LEFT | nCoV-2019_2 | GTTGATAAGTACTTTGATTGTTACGATGGT | 30 | 33.33 | 60.59 |
| nCoV-2019_50_RIGHT | nCoV-2019_2 | TAACATGTTGTGCCAACCACCA | 22 | 45.45 | 60.95 |
| nCoV-2019_51_LEFT | nCoV-2019_1 | TCAATAGCCGCCACTAGAGGAG | 22 | 54.55 | 61.34 |
| nCoV-2019_51_RIGHT | nCoV-2019_1 | AGTGCATTAACATTGGCCGTGA | 22 | 45.45 | 61.14 |
| nCoV-2019_52_LEFT | nCoV-2019_2 | CATCAGGAGATGCCACAACTGC | 22 | 54.55 | 61.83 |
| nCoV-2019_52_RIGHT | nCoV-2019_2 | GTTGAGAGCAAAATTCATGAGGTCC | 25 | 44.00 | 60.62 |
| nCoV-2019_53_LEFT | nCoV-2019_1 | AGCAAAATGTTGGACTGAGACTGA | 24 | 41.67 | 60.69 |
| nCoV-2019_53_RIGHT | nCoV-2019_1 | AGCCTCATAAAACTCAGGTTCCC | 23 | 47.83 | 60.31 |
| nCoV-2019_54_LEFT | nCoV-2019_2 | TGAGTTAACAGGACACATGTTAGACA | 26 | 38.46 | 60.18 |
| nCoV-2019_54_RIGHT | nCoV-2019_2 | AACCAAAAACTTGTCCATTAGCACA | 25 | 36.00 | 60.11 |
| nCoV-2019_55_LEFT | nCoV-2019_1 | ACTCAACTTTACTTAGGAGGTATGAGCT | 28 | 39.29 | 61.43 |
| nCoV-2019_55_RIGHT | nCoV-2019_1 | GGTGTACTCTCCTATTTGTACTTTACTGT | 29 | 37.93 | 60.54 |
| nCoV-2019_56_LEFT | nCoV-2019_2 | ACCTAGACCACCACTTAACCGA | 22 | 50.00 | 60.49 |
| nCoV-2019_56_RIGHT | nCoV-2019_2 | ACACTATGCGAGCAGAAGGGTA | 22 | 50.00 | 61.21 |
| nCoV-2019_57_LEFT | nCoV-2019_1 | ATTCTACACTCCAGGGACCACC | 22 | 54.55 | 61.16 |
| nCoV-2019_57_RIGHT | nCoV-2019_1 | GTAATTGAGCAGGGTCGCCAAT | 22 | 50.00 | 61.26 |
| nCoV-2019_58_LEFT | nCoV-2019_2 | TGATTTGAGTGTTGTCAATGCCAGA | 25 | 40.00 | 61.44 |
| nCoV-2019_58_RIGHT | nCoV-2019_2 | CTTTTCTCCAAGCAGGGTTACGT | 23 | 47.83 | 61.06 |
| nCoV-2019_59_LEFT | nCoV-2019_1 | TCACGCATGATGTTTCATCTGCA | 23 | 43.48 | 61.42 |
| nCoV-2019_59_RIGHT | nCoV-2019_1 | AAGAGTCCTGTTACATTTTCAGCTTG | 26 | 38.46 | 60.02 |
| nCoV-2019_60_LEFT | nCoV-2019_2 | TGATAGAGACCTTTATGACAAGTTGCA | 27 | 37.04 | 60.53 |
| nCoV-2019_60_RIGHT | nCoV-2019_2 | GGTACCAACAGCTTCTCTAGTAGC | 24 | 50.00 | 60.44 |
| nCoV-2019_61_LEFT | nCoV-2019_1 | TGTTTATCACCCGCGAAGAAGC | 22 | 50.00 | 61.50 |
| nCoV-2019_61_RIGHT | nCoV-2019_1 | ATCACATAGACAACAGGTGCGC | 22 | 50.00 | 61.25 |
| nCoV-2019_62_LEFT | nCoV-2019_2 | GGCACATGGCTTTGAGTTGACA | 22 | 50.00 | 61.91 |
| nCoV-2019_62_RIGHT | nCoV-2019_2 | GTTGAACCTTTCTACAAGCCGC | 22 | 50.00 | 60.35 |
| nCoV-2019_63_LEFT | nCoV-2019_1 | TGTTAAGCGTGTTGACTGGACT | 22 | 45.45 | 60.16 |
| nCoV-2019_63_RIGHT | nCoV-2019_1 | ACAAACTGCCACCATCACAACC | 22 | 50.00 | 61.85 |
| nCoV-2019_64_LEFT | nCoV-2019_2 | TCGATAGATATCCTGCTAATTCCATTGT | 28 | 35.71 | 60.11 |
| nCoV-2019_64_RIGHT | nCoV-2019_2 | AGTCTTGTAAAAGTGTTCCAGAGGT | 25 | 40.00 | 60.10 |
| nCoV-2019_65_LEFT | nCoV-2019_1 | GCTGGCTTTAGCTTGTGGGTTT | 22 | 50.00 | 61.92 |
| nCoV-2019_65_RIGHT | nCoV-2019_1 | TGTCAGTCATAGAACAAACACCAATAGT | 28 | 35.71 | 60.90 |
| nCoV-2019_66_LEFT | nCoV-2019_2 | GGGTGTGGACATTGCTGCTAAT | 22 | 50.00 | 61.21 |
| nCoV-2019_66_RIGHT | nCoV-2019_2 | TCAATTTCCATTTGACTCCTGGGT | 24 | 41.67 | 60.45 |
| nCoV-2019_67_LEFT | nCoV-2019_1 | GTTGTCCAACAATTACCTGAAACTTACT | 28 | 35.71 | 60.43 |
| nCoV-2019_67_RIGHT | nCoV-2019_1 | CAACCTTAGAAACTACAGATAAATCTTGGG | 30 | 36.67 | 60.40 |
| nCoV-2019_68_LEFT | nCoV-2019_2 | ACAGGTTCATCTAAGTGTGTGTGT | 24 | 41.67 | 60.14 |
| nCoV-2019_68_RIGHT | nCoV-2019_2 | CTCCTTTATCAGAACCAGCACCA | 23 | 47.83 | 60.31 |
| nCoV-2019_69_LEFT | nCoV-2019_1 | TGTCGCAAAATATACTCAACTGTGTCA | 27 | 37.04 | 61.43 |
| nCoV-2019_69_RIGHT | nCoV-2019_1 | TCTTTATAGCCACGGAACCTCCA | 23 | 47.83 | 61.14 |
| nCoV-2019_70_LEFT | nCoV-2019_2 | ACAAAAGAAAATGACTCTAAAGAGGGTTT | 29 | 31.03 | 60.13 |
| nCoV-2019_70_RIGHT | nCoV-2019_2 | TGACCTTCTTTTAAAGACATAACAGCAG | 28 | 35.71 | 60.27 |
| nCoV-2019_71_LEFT | nCoV-2019_1 | ACAAATCCAATTCAGTTGTCTTCCTATTC | 29 | 34.48 | 60.54 |
| nCoV-2019_71_RIGHT | nCoV-2019_1 | TGGAAAAGAAAGGTAAGAACAAGTCCT | 27 | 37.04 | 60.80 |
| nCoV-2019_72_LEFT | nCoV-2019_2 | ACACGTGGTGTTTATTACCCTGAC | 24 | 45.83 | 61.04 |
| nCoV-2019_72_RIGHT | nCoV-2019_2 | ACTCTGAACTCACTTTCCATCCAAC | 25 | 44.00 | 60.97 |
| nCoV-2019_73_LEFT | nCoV-2019_1 | CAATTTTGTAATGATCCATTTTTGGGTGT | 29 | 31.03 | 60.29 |
| nCoV-2019_73_RIGHT | nCoV-2019_1 | CACCAGCTGTCCAACCTGAAGA | 22 | 54.55 | 62.45 |
| nCoV-2019_74_LEFT | nCoV-2019_2 | ACATCACTAGGTTTCAAACTTTACTTGC | 28 | 35.71 | 60.68 |
| nCoV-2019_74_RIGHT | nCoV-2019_2 | GCAACACAGTTGCTGATTCTCTTC | 24 | 45.83 | 60.85 |
| nCoV-2019_75_LEFT | nCoV-2019_1 | AGAGTCCAACCAACAGAATCTATTGT | 26 | 38.46 | 60.24 |
| nCoV-2019_75_RIGHT | nCoV-2019_1 | ACCACCAACCTTAGAATCAAGATTGT | 26 | 38.46 | 60.69 |
| nCoV-2019_76_LEFT | nCoV-2019_2 | AGGGCAAACTGGAAAGATTGCT | 22 | 45.45 | 60.76 |
| nCoV-2019_76_RIGHT | nCoV-2019_2 | ACACCTGTGCCTGTTAAACCAT | 22 | 45.45 | 60.42 |
| nCoV-2019_77_LEFT | nCoV-2019_1 | CCAGCAACTGTTTGTGGACCTA | 22 | 50.00 | 60.75 |
| nCoV-2019_77_RIGHT | nCoV-2019_1 | CAGCCCCTATTAAACAGCCTGC | 22 | 54.55 | 61.59 |
| nCoV-2019_78_LEFT | nCoV-2019_2 | CAACTTACTCCTACTTGGCGTGT | 23 | 47.83 | 60.55 |
| nCoV-2019_78_RIGHT | nCoV-2019_2 | TGTGTACAAAAACTGCCATATTGCA | 25 | 36.00 | 60.22 |
| nCoV-2019_79_LEFT | nCoV-2019_1 | GTGGTGATTCAACTGAATGCAGC | 23 | 47.83 | 60.92 |
| nCoV-2019_79_RIGHT | nCoV-2019_1 | CATTTCATCTGTGAGCAAAGGTGG | 24 | 45.83 | 60.62 |
| nCoV-2019_80_LEFT | nCoV-2019_2 | TTGCCTTGGTGATATTGCTGCT | 22 | 45.45 | 60.89 |
| nCoV-2019_80_RIGHT | nCoV-2019_2 | TGGAGCTAAGTTGTTTAACAAGCG | 24 | 41.67 | 60.02 |
| nCoV-2019_81_LEFT | nCoV-2019_1 | GCACTTGGAAAACTTCAAGATGTGG | 25 | 44.00 | 61.24 |
| nCoV-2019_81_RIGHT | nCoV-2019_1 | GTGAAGTTCTTTTCTTGTGCAGGG | 24 | 45.83 | 60.73 |
| nCoV-2019_82_LEFT | nCoV-2019_2 | GGGCTATCATCTTATGTCCTTCCCT | 25 | 48.00 | 61.52 |
| nCoV-2019_82_RIGHT | nCoV-2019_2 | TGCCAGAGATGTCACCTAAATCAA | 24 | 41.67 | 60.02 |
| nCoV-2019_83_LEFT | nCoV-2019_1 | TCCTTTGCAACCTGAATTAGACTCA | 25 | 40.00 | 60.46 |
| nCoV-2019_83_RIGHT | nCoV-2019_1 | TTTGACTCCTTTGAGCACTGGC | 22 | 50.00 | 61.33 |
| nCoV-2019_84_LEFT | nCoV-2019_2 | TGCTGTAGTTGTCTCAAGGGCT | 22 | 50.00 | 61.61 |
| nCoV-2019_84_RIGHT | nCoV-2019_2 | AGGTGTGAGTAAACTGTTACAAACAAC | 27 | 37.04 | 60.36 |
| nCoV-2019_85_LEFT | nCoV-2019_1 | ACTAGCACTCTCCAAGGGTGTT | 22 | 50.00 | 61.03 |
| nCoV-2019_85_RIGHT | nCoV-2019_1 | ACACAGTCTTTTACTCCAGATTCCC | 25 | 44.00 | 60.51 |
| nCoV-2019_86_LEFT | nCoV-2019_2 | TCAGGTGATGGCACAACAAGTC | 22 | 50.00 | 61.07 |
| nCoV-2019_86_RIGHT | nCoV-2019_2 | ACGAAAGCAAGAAAAAGAAGTACGC | 25 | 40.00 | 61.01 |
| nCoV-2019_87_LEFT | nCoV-2019_1 | CGACTACTAGCGTGCCTTTGTA | 22 | 50.00 | 60.16 |
| nCoV-2019_87_RIGHT | nCoV-2019_1 | ACTAGGTTCCATTGTTCAAGGAGC | 24 | 45.83 | 60.81 |
| nCoV-2019_88_LEFT | nCoV-2019_2 | CCATGGCAGATTCCAACGGTAC | 22 | 54.55 | 61.58 |
| nCoV-2019_88_RIGHT | nCoV-2019_2 | TGGTCAGAATAGTGCCATGGAGT | 23 | 47.83 | 61.40 |
| nCoV-2019_89_LEFT | nCoV-2019_1 | GTACGCGTTCCATGTGGTCATT | 22 | 50.00 | 61.50 |
| nCoV-2019_89_RIGHT | nCoV-2019_1 | ACCTGAAAGTCAACGAGATGAAACA | 25 | 40.00 | 60.91 |
| nCoV-2019_90_LEFT | nCoV-2019_2 | ACACAGACCATTCCAGTAGCAGT | 23 | 47.83 | 61.58 |
| nCoV-2019_90_RIGHT | nCoV-2019_2 | TGAAATGGTGAATTGCCCTCGT | 22 | 45.45 | 60.82 |
| nCoV-2019_91_LEFT | nCoV-2019_1 | TCACTACCAAGAGTGTGTTAGAGGT | 25 | 44.00 | 60.93 |
| nCoV-2019_91_RIGHT | nCoV-2019_1 | TTCAAGTGAGAACCAAAAGATAATAAGCA | 29 | 31.03 | 60.03 |
| nCoV-2019_92_LEFT | nCoV-2019_2 | TTTGTGCTTTTTAGCCTTTCTGCT | 24 | 37.50 | 60.14 |
| nCoV-2019_92_RIGHT | nCoV-2019_2 | AGGTTCCTGGCAATTAATTGTAAAAGG | 27 | 37.04 | 60.53 |
| nCoV-2019_93_LEFT | nCoV-2019_1 | TGAGGCTGGTTCTAAATCACCCA | 23 | 47.83 | 61.59 |
| nCoV-2019_93_RIGHT | nCoV-2019_1 | AGGTCTTCCTTGCCATGTTGAG | 22 | 50.00 | 60.55 |
| nCoV-2019_94_LEFT | nCoV-2019_2 | GGCCCCAAGGTTTACCCAATAA | 22 | 50.00 | 60.56 |
| nCoV-2019_94_RIGHT | nCoV-2019_2 | TTTGGCAATGTTGTTCCTTGAGG | 23 | 43.48 | 60.18 |
| nCoV-2019_95_LEFT | nCoV-2019_1 | TGAGGGAGCCTTGAATACACCA | 22 | 50.00 | 61.10 |
| nCoV-2019_95_RIGHT | nCoV-2019_1 | CAGTACGTTTTTGCCGAGGCTT | 22 | 50.00 | 61.95 |
| nCoV-2019_96_LEFT | nCoV-2019_2 | GCCAACAACAACAAGGCCAAAC | 22 | 50.00 | 61.82 |
| nCoV-2019_96_RIGHT | nCoV-2019_2 | TAGGCTCTGTTGGTGGGAATGT | 22 | 50.00 | 61.36 |
| nCoV-2019_97_LEFT | nCoV-2019_1 | TGGATGACAAAGATCCAAATTTCAAAGA | 28 | 32.14 | 60.22 |
| nCoV-2019_97_RIGHT | nCoV-2019_1 | ACACACTGATTAAAGATTGCTATGTGAG | 28 | 35.71 | 60.17 |
| nCoV-2019_98_LEFT | nCoV-2019_2 | AACAATTGCAACAATCCATGAGCA | 24 | 37.50 | 60.50 |
| nCoV-2019_98_RIGHT | nCoV-2019_2 | TTCTCCTAAGAAGCTATTAAAATCACATGG | 30 | 33.33 | 60.01 |
